# RNF13 Regulates the Endolysosomal Pathway Through Interaction with the Small GTPase Arl8B

**DOI:** 10.1101/2025.07.31.667977

**Authors:** Audrey M. Sénécal, Valérie C. Cabana, Antoine Y. Bouchard, Laurent Cappadocia, Marc P. Lussier

## Abstract

The endolysosomal system is a dynamic intracellular network essential for cargo degradation, recycling and spatial compartmentalization. Precise coordination of endosome maturation and positioning is critical for maintaining lysosomal function and regulating receptor fate. This study uncovers a novel role for the E3 ubiquitin ligase RNF13 in controlling endolysosomal dynamics through its interaction with the small GTPase Arl8B. Using predictive structural modeling and co-immunoprecipitation assays, the results demonstrate that RNF13 binds to Arl8B, implicating residues Glu22 and Phe55 of Arl8B with RNF13’s Leu244. Their interaction influences lysosomal positioning and the trafficking of endocytic cargo. Notably, loss of RNF13-Arl8B binding alters Arl8B localization and causes a peripheral redistribution of lysosomes, while not affecting the abundance of endolysosomal markers. However, it does impair the internalization of the epidermal growth factor receptor (EGFR). These findings suggest that the RNF13-Arl8B interaction plays a crucial role in modulating vesicle maturation and fusion. Furthermore, overexpression of the Arl8B effector PLEKHM1 enhances RNF13-Arl8B complex formation, indicating a possible cooperative assembly of tethering complexes during lysosome– endosome fusion. Together, the results identify RNF13 as a spatial regulator of lysosomal organization and cargo processing, operating through a non-enzymatic scaffolding mechanism. This reveals an additional layer of regulation in endolysosomal trafficking, supporting a role for RNF13 as a checkpoint in cargo progression through degradative pathways. Altogether, the results of this work expand the understanding of the molecular coordination underlying lysosomal dynamics and underscore the importance of selective effector interactions in coordinating endolysosomal trafficking.

## INTRODUCTION

The endolysosomal pathway is an essential cellular network responsible for regulating intracellular trafficking, degradation and recycling of macromolecules, thereby maintaining cellular homeostasis and supporting physiological processes such as receptor signaling [1, 2]. This pathway is characterized by a series of dynamic and functionally distinct compartments, including early endosomes, late endosomes and lysosomes, which must mature, move and fuse in a tightly coordinated manner to ensure efficient cargo processing and proper spatial organization within the cell [3, 4]. Spatial positioning of endolysosomal compartments has emerged as a critical regulatory feature of their function. Lysosomes, in particular, can localize anywhere from the cell periphery to the perinuclear region, and their distribution directly affects their degradative capacity and fusion efficiency with other vesicles [5, 6]. This positioning is governed by small GTPases of the Ras-related protein Rab and ARL (ADP-ribosylation factor-like) families, which operate through specific effector proteins to recruit cytoskeletal motor complexes [7–9].

The small GTPase ADP-ribosylation factor-like 8B (Arl8B) has emerged as a key regulator of lysosomal motility and positioning, by orchestrating both anterograde and retrograde lysosomal transport by engaging with distinct sets of effectors [5, 10, 11]. It recruits SKIP (also known as PLEKHM2) to drive kinesin-mediated movement toward the cell periphery while interacting with PLEKHM1 to promote HOPS complex assembly and lysosome-endosome fusion [10, 12, 13]. Conversely, Arl8B can associate with RUFY3 and RUFY4 to mediate retrograde dynein-dynactin transport [14]. These bidirectional interactions are essential for the dynamic control of lysosomal positioning in response to cellular needs [15, 16]. The capacity of Arl8B to perform its function is tightly linked to its nucleotide-bound state. Like other small GTPases, Arl8B alternates between an inactive GDP-bound and an active GTP-bound form, with the GTP-loaded state being necessary for effector engagement and downstream trafficking activity [17, 18]. For instance, the GTP-bound form of Arl8B is essential for its interaction with various effectors, such as the RUN domains of PLEKHM1 and SKIP, both of which are involved in lysosomal fusion and transport [19]. On the other hand, GDP-bound Arl8B was demonstrated to localize to lipid droplets and facilitate the formation of lipid-droplet-lysosome contacts [20].

Among the known regulators of Arl8B is the RING finger protein 167 (RNF167), a PA-TM-RING family E3 ubiquitin ligase [21]. RNF167 has been shown to directly ubiquitinate Arl8B, targeting it for proteasomal degradation and thereby influencing lysosomal positioning and endocytic trafficking efficiency [22]. Specifically, RNF167-mediated Arl8B degradation results in the repositioning of lysosomes toward the perinuclear region and the suppression of peripheral trafficking events, indicating a degradation-based mechanism of spatial control.

In parallel, increasing attention is being paid to the role of E3 ubiquitin ligases (UbE3) in regulating vesicular trafficking beyond their classical function in protein degradation [23–26]. One such UbE3 is the RING finger protein 13 (RNF13), another member of the PA-TM-RING family that shares domain architecture and subcellular localization features with RNF167, which localizes to the endoplasmic reticulum (ER), late endosomes, and lysosomes [27].

RNF13 has been implicated in various cellular processes, including protein degradation, trafficking, and localization [27–29]. It has been shown to regulate the ubiquitination of the SNARE complex-associated protein SNAPIN (also known as BORCS3), a component of the BORC complex involved in Arl8B recruitment to lysosomes and to control the turnover of the lysosomal membrane protein Lamp1 [28–30]. More specifically, RNF13 was shown to promote the proteosomal degradation of Lamp1, thereby impairing lysosomal integrity and suppressing the degradation of Toll-like receptor (TLR) in lysosomes. This degradation of Lamp1 was also linked to impaired autophagic flux, suggesting that RNF13 can modulate both innate immune signaling and autophagy through its control of lysosomal membrane components [29].

Given the overlapping localization and structural homology between RNF13 and RNF167 [21], it is plausible that RNF13 might exert similar or complementary effects on Arl8B-dependent trafficking. Supporting this notion, a recent study has shown that RNF13 can modulate Arl8B ubiquitination in a pH- and Ca^2+^-dependent manner [31]. In this study, we focus on the physical and functional interaction between RNF13 and Arl8B, independently of RNF13’s ubiquitin ligase activity. Notably, although typically defined by their enzymatic roles in ubiquitination, several E3 ubiquitin ligases have been implicated in non-catalytic functions, acting as scaffolds or spatial regulators within trafficking pathways [32]. This broader functional versatility raises the possibility that RNF13, given its localization and known associations with lysosomal components, may also play a structural role in coordinating endolysosomal dynamics. Yet, the exact molecular mechanisms by which RNF13 contributes to trafficking dynamics remain poorly understood.

In this study, we identified Arl8B as a functionally relevant interactor of RNF13. Using structure-guided interaction prediction and co-immunoprecipitation assays, we demonstrate that RNF13 binds to Arl8B via conserved residues Glu22 and Phe55, sites previously identified as essential for the binding of Arl8B effectors PLEKHM1 and SKIP [19]. However, unlike canonical Arl8B effectors that preferentially engage the GTP-bound form, we found that RNF13 interacts with Arl8B regardless of its nucleotide-bound state. Alphafold3 structural modeling predicted stable RNF13–Arl8B interfaces in both GTP- and GDP-bound conformations, and co-immunoprecipitation assays confirmed that RNF13 binds both constitutively active (Q75L) and inactive (T34N) Arl8B variants. Furthermore, this interaction is critical for RNF13’s lysosomal localization and for maintaining perinuclear clustering of lysosomes. Disruption of this interaction, either by mutating Arl8B binding residues or depleting RNF13, causes lysosomes and late endosomes to redistribute toward the cell periphery, consistent with impaired retrograde trafficking [5, 14]. We further show that the RNF13-Arl8B interaction impacts lysosomal cargo processing. Indeed, cells expressing interaction-defective mutants of either RNF13 or Arl8B exhibit impaired EGFR internalization kinetics, suggesting that their interaction modulates vesicle maturation or fusion events. Interestingly, co-expression of PLEKHM1 enhances RNF13-Arl8B complex formation, raising the possibility that multiple Arl8B effectors may interact in a coordinated or stepwise manner at the lysosome surface to regulate membrane tethering and fusion events. Together, our results reveal a previously uncharacterized role for RNF13 in controlling endolysosomal architecture and cargo degradation. Moreover, the ability of RNF13 to bind Arl8B independently of its activation state supports a model in which RNF13 contributes to the regulation of Arl8B-dependent trafficking events throughout its nucleotide cycle, revealing a new layer of regulatory complexity in endolysosomal dynamics.

## RESULTS

### RNF13 associates with Arl8B in a cellular context

Given our laboratory’s ongoing interest in the functional roles of PA-TM-RING E3 ubiquitin ligases, RNF167 and RNF13, we sought to investigate whether RNF13 may influence the endolysosomal system in ways similar to those previously established for its homolog RNF167 [21, 27]. Notably, RNF167 has been shown to regulate lysosomal positioning and endocytic trafficking by targeting the small GTPase Arl8B for proteasomal degradation [22]. This finding prompted us to consider Arl8B as a possible interactor for RNF13, especially considering their shared subcellular localization. As emerging evidence demonstrates RNF13 involvement in vesicular trafficking and lysosomal regulation [21, 28, 29, 31], we explored whether RNF13 might also functionally intersect with Arl8B-dependent pathways. To strengthen this rationale and guide our experimental approach, we first examined the structural feasibility of such an interaction. Using structure-guided interaction modeling with AlphaFold, we predicted a potential binding interface between RNF13 and Arl8B. The resulting structural model revealed a possible interaction between Arl8B and the RING domain of RNF13, a region associated with its catalytic ubiquitin ligase activity (Fig. 1A). This prediction supported the hypothesis that RNF13 could associate with Arl8B, motivating the exploration of this interaction in a cellular context.

**Figure 1:**
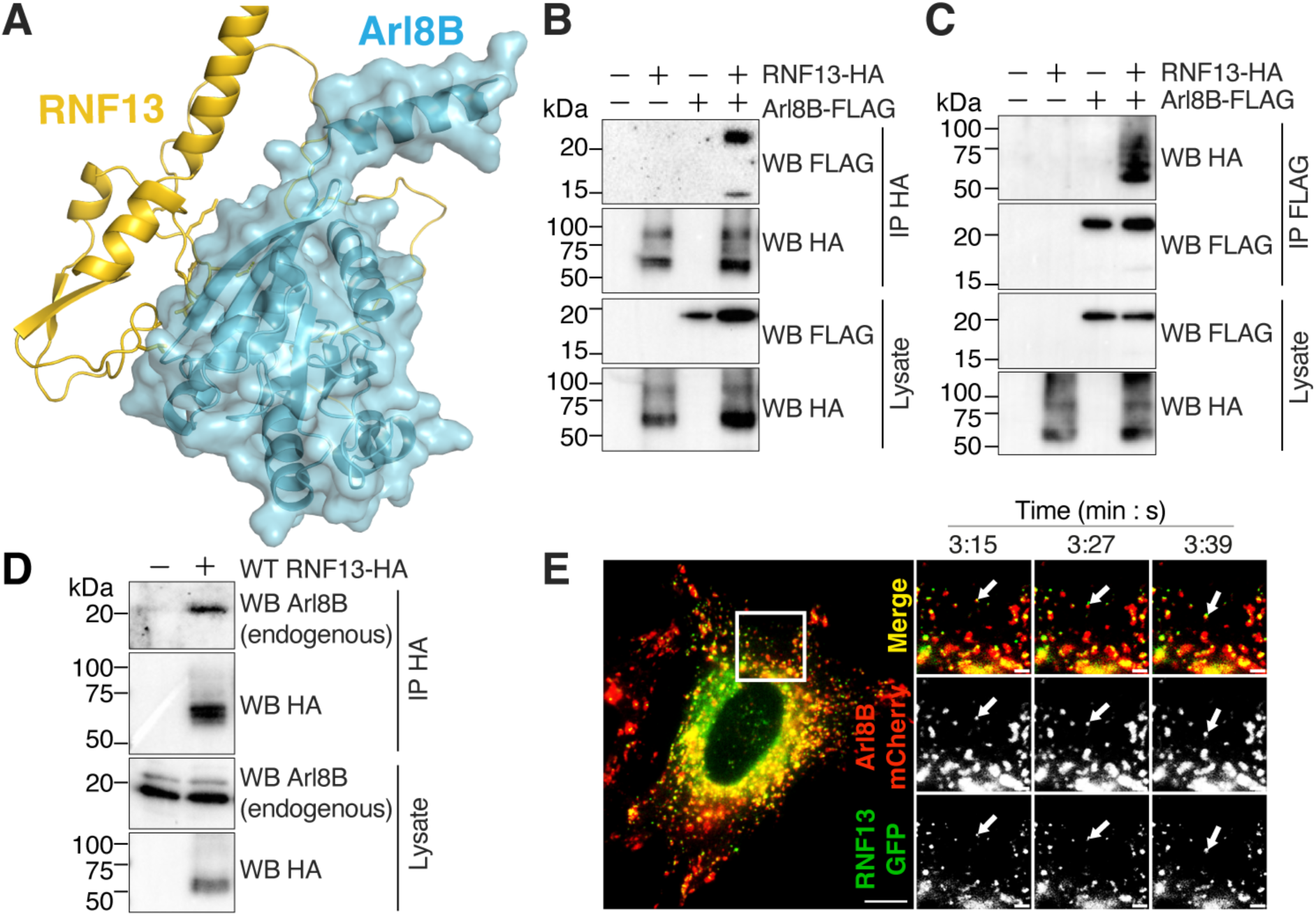
RNF13 interacts with Arl8B. (**A**) Predicted complex of Arl8B (blue) and RNF13 (yellow), generated using ColabFold [48, 49]. Key interacting residues of RNF13 within the RING domain are shown as sticks, highlighting the predicted interface with Arl8B. (**B-D**) Western blots of co-immunoprecipitation assays in transfected HEK293T/17 cells show binding between Arl8B (FLAG-tagged in **B-C**; endogenous Arl8B in **D**) and RNF13-HA. (**E**) Representative live-cell images of HeLa cells expressing RNF13-GFP (in green) and Arl8B-mCherry (in red). White arrowheads represent an event where RNF13-GFP and Arl8B-mCherry initially overlap, then slightly dissociate and move together. Results are representative of three independent experiments (N=3).

Using co-immunoprecipitation assays, we confirmed that wild-type (WT) Arl8B interacts with WT RNF13 in a cellular context (Fig. 1B-D). This result was consistently observed regardless of whether RNF13 (Fig. 1B) or Arl8B (Fig. 1C) was used as the bait. Still, more importantly, it could also be detected with endogenous Arl8B following RNF13-HA immunoprecipitation (Fig. 1D). Live-cell epifluorescence imaging was conducted on transiently transfected HeLa cells to gain insight into the dynamics between RNF13-GFP and Arl8B-mCherry. The results are consistent with the formation of a transient RNF13-Arl8B complex, part of an overall coordinated movement (Fig. 1E, SMovie 1), which subsequently dissociates (SMovie 2). Notably, RNF13-Arl8B complexes were almost immobile in the perinuclear region compared to their greater motility at the cell periphery, suggesting spatially distinct interaction dynamics. RNF13-GFP displayed a broad distribution, with substantial accumulation in the perinuclear area and additional signal throughout the cytoplasm, exhibiting a dynamic movement consistent with vesicle trafficking (Fig. 1E). In contrast, Arl8B-mCherry showed a more confined pattern, characterized by highly motile puncta that accumulated at the cell periphery and exhibited a noticeably weaker signal in the perinuclear region (Fig. 1E). While areas of overlap between RNF13 and Arl8B were observed, especially on moving vesicles, these spatial distinctions suggest that they partially occupy separate vesicle populations, with their interaction occurring transiently on dynamic vesicle. Overall, these results provide evidence that RNF13 and Arl8B can form complexes and interact with each other.

### RNF13 and Arl8B have distinct yet intersecting roles in lysosomal and endosomal distribution

To assess how the depletion of either RNF13 or Arl8B impacts the other, as well as the endolysosomal pathway, we used RNA interference. The efficiency of DsiRNAs (Dicer-substrate small interfering RNAs) targeting endogenous Arl8B was first validated by immunoblotting, which confirmed a robust knockdown (Fig. 2A-B). As previously demonstrated [33], Arl8B depletion did not affect the distribution of early endosomes, as shown by the unchanged localization of EEA1 (Fig. 2C, J). However, Arl8B knockdown led to an increased colocalization of RNF13 with early endosomal marker EEA1 (Fig. 2C-D) and late endosomal marker Rab7 (Fig. 2E-F), indicating a redistribution of RNF13 towards these compartments. In contrast, its colocalization with Lamp1-positive lysosomes was reduced (Fig. 2G-H), despite RNF13’s overall distribution remaining unchanged (Fig. 2I). These results suggest that Arl8B is required for efficient targeting of RNF13 to lysosomes. To further explore the role of Arl8B in organelle positioning, we examined whether its knockdown affected the spatial distribution of endolysosomal compartments. Notably, neither Arl8B knockdown nor RNF13 overexpression alone significantly altered the spatial distribution of Rab7 (Fig. 2K). However, when Arl8B knockdown and RNF13 overexpression were combined, a significant increase in the peripheral localization of Rab7-positive vesicles was observed (Fig. 2E, K), suggesting that RNF13 may influence late endosomal positioning in the absence of Arl8B. Similarly and consistent with Arl8B’s established role in promoting peripheral lysosome localization [17], its depletion with siArl8B#2 in particular, led to a reduction in the peripheral distribution of Lamp1-positive vesicles (Fig. 2G, L). Interestingly, while RNF13 overexpression alone did not significantly alter lysosome distribution, its combination with Arl8B’s depletion significantly enhanced the perinuclear clustering of lysosomes (Fig. 2L), suggesting an additive effect that reinforces lysosome retention near the nucleus in the absence of Arl8B. Together, these results reveal a reciprocal relationship between RNF13 and Arl8B, where Arl8B is essential for directing RNF13 to lysosomes. At the same time, RNF13 modulates the spatial organization of late endosomes and lysosomes in conditions when Arl8B is depleted.

**Figure 2:**
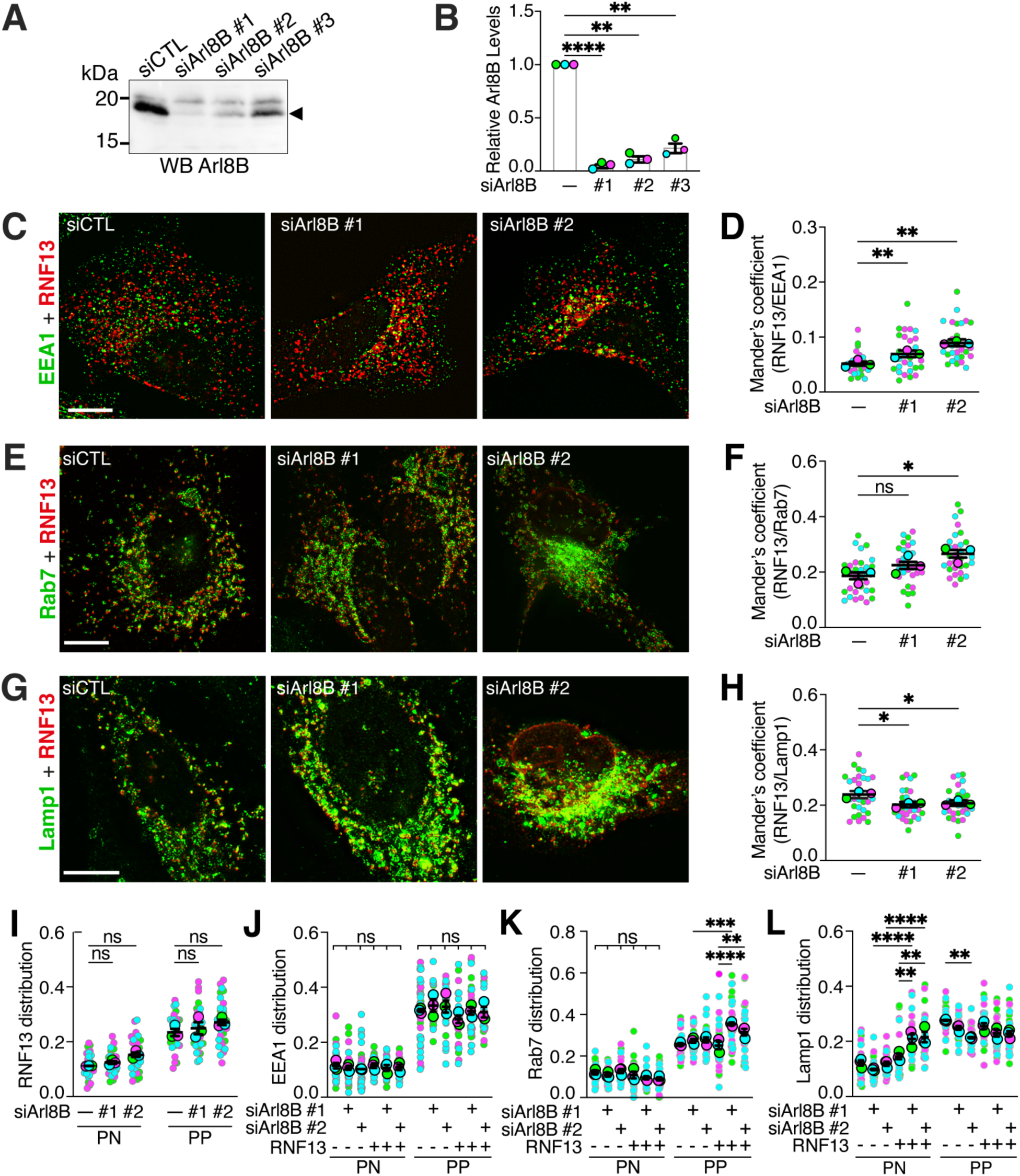
Arl8B knockdown affects the subcellular distribution of endolysosomal compartments. (**A**) Representative immunoblot shows endogenous Arl8B detection from the lysate of HEK293T/17 cells transfected with 10 nM of Dicer-substrate siRNAs (DsiRNAs) targeting Arl8B or a non-targeting control (siCTL). Band densities were analyzed and normalized to the stain-free signal (loading control) and then to the siCTL lane using ImageLab software. (**B**) The strip plots show the average relative protein abundance of Arl8B detected from three independent experiments (N=3), with each experiment represented by a distinct color. (**C, E, G**) Representative immunofluorescence images show HeLa cells transfected with dsiRNAs targeting Arl8B, stained for RNF13-HA (in red) and specific compartment markers (in green): (**C**) early endosomes (EEA1), (**E**) late endosomes (Rab7) and (**G**) lysosomes (Lamp1). Scale bars indicate 10 µm. (**D, F, H-L**) The jittered individual value plots represent (**D, F, H**) Mander’s overlap coefficient of RNF13-HA with indicated markers in Arl8B KD cells or (**I-L**) the spatial distribution of the respective endolysosomal marker (EEA1, Rab7, Lamp1) and RNF13-HA. Each color represents an independent experiment (N=3), with 10 cells analyzed per experiment (n=30). Larger circles indicate the experimental mean while smaller circles represent individual cell measurements. All results are expressed as mean ± SEM from N=3. Statistical significance was assessed using (**B, D, F, H**) one-way or (**I-L**) two-way RM-ANOVA with multiple comparisons test. ns: not significant; * *p* < 0.05; ** *p* < 0.01; *** *p* < 0.001; **** *p* < 0.0001; PN = perinuclear; PP = peripheral.

According to recent literature, RNF13 influences the size of endolysosomal vesicles [28] and plays a role in lysosomal regulation [29]. Thus, we also investigated how RNF13 depletion affects the distribution of late endosomes and lysosomes. The effectiveness of RNF13 knockdown was confirmed by immunoblotting, which showed a substantial reduction in the abundance of endogenous RNF13 protein following treatment with specific DsiRNAs (Fig. 3A-B). Interestingly, RNF13 knockdown did not significantly impact Arl8B’s colocalization with either Rab7- or Lamp1-positive compartments (Fig. 3C-F), suggesting that RNF13 does not directly regulate Arl8B’s recruitment to late endosomes or lysosomes. Consistent with recent findings implicating RNF13 in promoting perinuclear lysosomal positioning [31], Arl8B exhibited a significant change toward the cell periphery in RNF13-depleted cells (Fig. 3G). On the other hand, RNF13 knockdown alone did not significantly alter the spatial distribution of late endosomes or lysosomes (Fig. 3H-I). However, the co-expression of exogenous Arl8B in RNF13-depleted cells resulted in a significant increase in the peripheral localization of Rab7- positive late endosomes (Fig. 3C, H). Notably, Arl8B overexpression alone (in both siCTL conditions) was associated with a modest but consistent increase in the perinuclear localization of Rab7 without impacting its peripheral distribution (Fig. 3H). These findings were unexpected, as Arl8B overexpression is typically associated with Rab7 displacement from peripheral lysosomal compartments via the SKIP-HOPS-TBC1D15 pathway [34]. The observed increase in peripheral Rab7 under RNF13 depletion suggests that RNF13 may usually contribute to regulating the spatial distribution of late endosomal compartments by limiting Rab7 retention at the cell periphery. Similarly, Arl8B overexpression alone led to a general increase in the peripheral localization of Lamp1-positive lysosomes, consistent with its known role in promoting anterograde lysosomal transport [33]. Still, this redistribution was further amplified when RNF13 was depleted, suggesting an additive effect on lysosome positioning (Fig. 3I). Altogether, these results indicate that RNF13 acts as a modulator of endolysosomal spatial organization, constraining the peripheral redistribution of both Rab7- and Lamp1-positive compartments, particularly in the context of elevated Arl8B activity.

**Figure 3:**
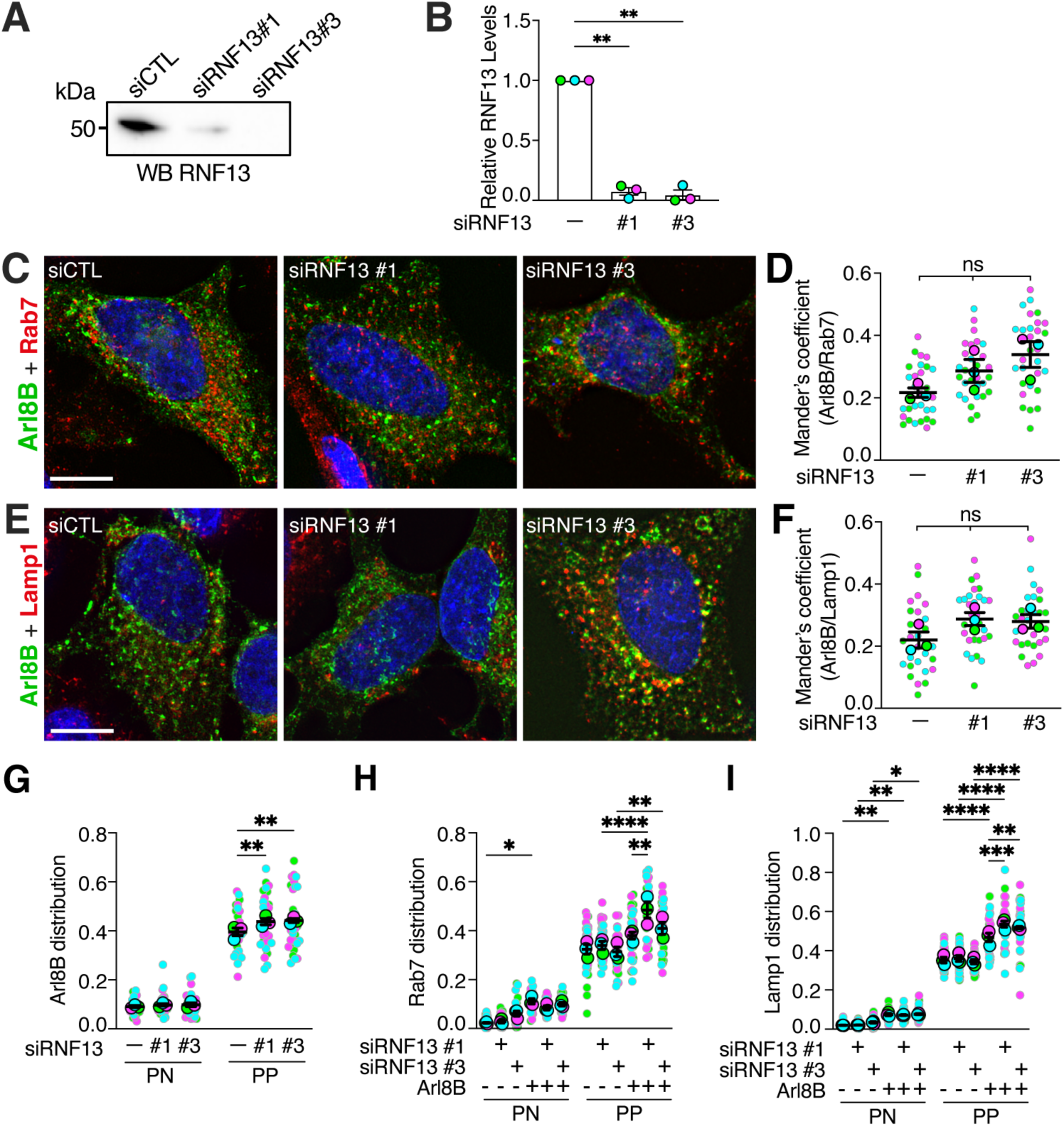
RNF13 depletion enhances Arl8B-driven peripheral redistribution of late endosomes and lysosomes. (**A**) Representative immunoblot shows endogenous RNF13 detection from the lysate of HEK293T/17 cells transfected with 10 nM of Dicer-substrate siRNAs (DsiRNAs) targeting RNF13 or a non-targeting control (siCTL). Band densities were analysed as previously described. (**B**) The strip plots show the average relative protein abundance of endogenous RNF13 detected from three independent experiments (N=3), with each experiment represented by a distinct color. (**C, E**) Representative immunofluorescence images show HeLa cells transfected with siRNF13, stained for Arl8B-FLAG (in green), along with markers for distinct endosomal compartments (in red): (**C**) late endosomes (Rab7) and (**E**) lysosomes (Lamp1). Scale bars indicate 10 µm. (**D, F, G-I**) The jittered individual value plots represent (**D, F**) Mander’s overlap coefficient of Arl8B-FLAG with indicated markers in RNF13 KD cells or (**G-I**) the spatial distribution of the respective endolysosomal marker (Rab7, Lamp1) and Arl8B-FLAG. Each color represents an independent experiment (N=3), with 10 cells analyzed per experiment (n=30). Larger circles indicate the experimental mean, and the smaller circles represent individual cell measurements. All results are expressed as mean ± SEM from N=3. Statistical significance was assessed using (**B, D, F**) one-way or (**G-I**) two-way RM-ANOVA with multiple comparisons test. ns: not significant; * *p* < 0.05; ** *p* < 0.01; *** *p* < 0.001; **** *p* < 0.0001; PN = perinuclear; PP = peripheral.

### Arl8B residues Glu22 and Phe55 are essential for RNF13 binding and influence endosomal compartment association

We next sought to elucidate the molecular basis of RNF13-Arl8B interaction. AlphaFold’s predicted structure revealed key residues potentially stabilizing the RNF13 and Arl8B complex. The predicted structure notably reveals a hydrogen bond between RNF13 Lys282 and Arl8B Glu22 alongside the embedding of RNF13 Leu244 in a hydrophobic pocket of Arl8B formed by Val53, Phe78, Met81, and Phe55. Among these residues, Arl8B Phe55 was identified as the principal contact site (Fig. 4A). To experimentally validate this binding model, we performed co-immunoprecipitation assays. Mutation of Arl8B residues Glu22 and Phe55 abolished co-immunoprecipitation with RNF13, indicating an important role in forming the complex (Fig. 4B). Results from immunofluorescence microscopy further showed a reduced colocalization of Arl8B variants with RNF13, supporting the structural prediction (Fig. 4C-D).

**Figure 4:**
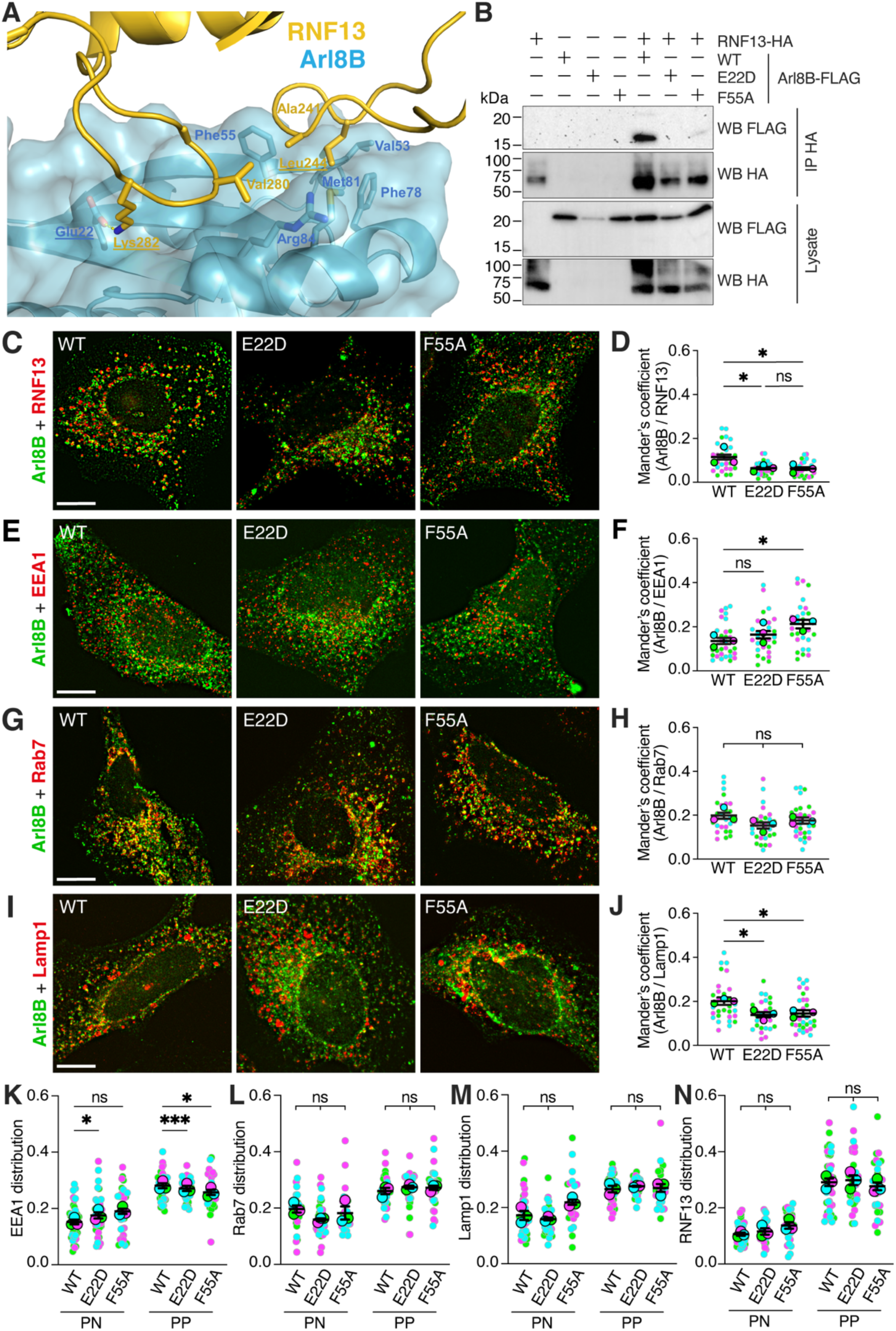
Mutations at Glu22 or Phe55 of Arl8B disrupt RNF13-Arl8B interaction. (**A**) Predicted complex of Arl8B (blue) and RNF13 (yellow) showing their interacting residues, generated using ColabFold [48, 49]. (**B**) Representative (N=3) immunoblots of co-immunoprecipitation from HEK293T/17 cells transiently expressing FLAG-tagged Arl8B (WT or variants E22D, F55A) and HA-tagged RNF13. (**C**) Representative (N=3) immunofluorescence images of RNF13 (red) with Arl8B (WT, E22D or F55A; green) and (**D**) their corresponding Mander’s overlap coefficient jitter plot. (**E, G, I**) Representative immunofluorescence images showing Arl8B-FLAG (WT, E22D or F55A; in green) with endolysosomal markers (in red): (**E**) EEA1, (**G**) Rab7 and (**I**) Lamp1. Scale bars indicate 10 µm. (**F, H, J-N**) The jittered individual value plots represent (**F, H, J**) Mander’s overlap coefficient of Arl8B with indicated markers or (**K-N**) the spatial distribution of (**K-M**) the specific compartment markers and (**N**) RNF13-HA. Each color represents an independent experiment (N=3), with 10 cells analyzed per experiment (n=30). Larger circles indicate the experimental mean while smaller circles represent individual cell measurements. All results are expressed as mean ± SEM from N=3. Statistical significance was assessed using (**D, F, H, J**) one-way or (**K-N**) two-way RM-ANOVA with multiple comparisons test. ns: not significant; * *p* < 0.05; *** *p* < 0.001; PN = perinuclear; PP = peripheral.

We next investigated the consequences of substituting residues Glu22 and Phe55 on Arl8B’s localization. Although RNF13 knockdown did not affect the overall localization of Arl8B (Fig. 3C-F), Arl8B E22D and F55A variants significantly altered its subcellular pattern, as reflected by changes in its colocalization with distinct endolysosomal compartments. Specifically, these variants exhibited increased colocalization with EEA1-positive early endosomes (Fig. 4E-F) while their presence on Rab7-positive late endosomes remained unchanged (Fig. 4G-H). In contrast, their colocalization with Lamp1-positive lysosomes was significantly reduced (Fig. 4I-J). Interestingly, expression of these binding-defective variants did not significantly alter the distribution of RNF13, late endosomes or lysosomes (Fig. 4L-N). However, a subtle yet significant redistribution of early endosomes towards the perinuclear region was measured (Fig. 4K). These results establish Glu22 and Phe55 of Arl8B as key structural determinants for binding RNF13 and demonstrate that disruption of this interface redistributes Arl8B away from lysosomes and toward early endosomal compartments.

### RNF13 residue Leu244 is required for Arl8B interaction and contributes to the positioning of early endosomes and lysosomes

Following our structural predictions that implicated Leu244 in stabilizing the RNF13 interaction with the Arl8B hydrophobic pocket (Fig. 4A), we validated the functional significance of this residue. Co-immunoprecipitation assays demonstrated that substituting Leu244 with an alanine (L244A) substantially impaired RNF13’s ability to bind WT Arl8B (Fig. 5A). This disruption was further supported by immunofluorescence analysis, which revealed a marked decrease in colocalization between RNF13 L244A and WT Arl8B when compared to RNF13 WT (Fig. 5B-C), reinforcing the essential role of Leu244 in facilitating the interaction. Next, we examined how the L244A substitution affects RNF13’s localization. The L244A variant resulted in a significant increase in colocalization with EEA1-positive early endosomes compared to RNF13 WT (Fig. 5D-E). In contrast, RNF13 colocalization with late endosome marker Rab7 and lysosomal marker Lamp1 remained unaffected (Fig. 5F-I). However, a modest, non-significant reduction in RNF13 L244A variant colocalization with Lamp1 was observed, suggesting that Arl8B binding via Leu244 may contribute to, but is not strictly required for RNF13’s recruitment to late compartments.

**Figure 5:**
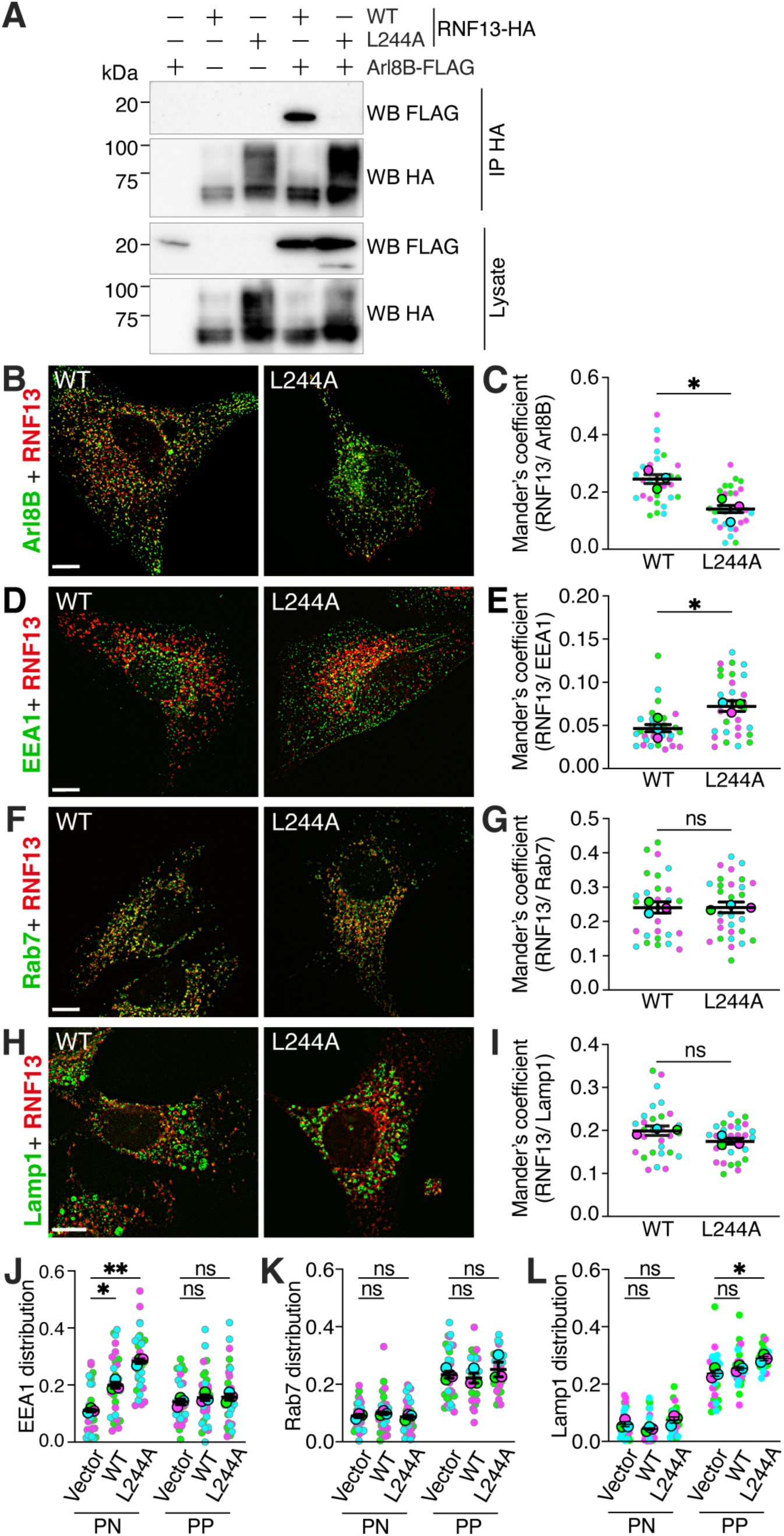
Mutation at Leu244 of RNF13 is enough to destabilize its interaction with Arl8B. (**A**) Representative (N=3) immunoblots of co-immunoprecipitation from HEK293T/17 cells transiently expressing HA-tagged RNF13 (WT or L244A) and FLAG-tagged Arl8B. (**B, D, F, H**) Representative (N=3) immunofluorescence images of RNF13 WT or L244A (in red) with (**B**) Arl8B (in green) or endogenous endolysosomal marker (in green) (**D**) EEA1, (**F**) Rab7 and (**H**) Lamp1. Scale bars indicate 10 µm. (**C, E, G, I-L**) The jittered individual value plots represent (**C, E, G, I**) Mander’s overlap coefficient of RNF13 with indicated markers or (**J-L**) the spatial distribution of the specific compartment markers. Each color represents an independent experiment (N=3), with 10 cells analyzed per experiment (n=30). Larger circles indicate the experimental mean while smaller circles represent individual cell measurements. All results are expressed as mean ± SEM from N=3. Statistical significance was assessed using (**C, E, G, I**) one-way or (**J-L**) two-way RM-ANOVA with multiple comparisons test. ns: not significant; * *p* < 0.05; ** *p* < 0.01; PN = perinuclear; PP = peripheral.

To assess whether the RNF13 L244A variant affects endosomal positioning, we analyzed the spatial distribution of endosomal compartments. Ectopic expression of RNF13 WT led to a perinuclear accumulation of EEA1-positive early endosomes, which was further enhanced by the expression of RNF13 L244A variant (Fig. 5J). This contrasts with the unchanged EEA1 distribution observed upon RNF13 WT overexpression in siCTL-transfected cells (Fig. 2J), possibly reflecting the impact of lipid-based siRNA delivery systems on native endosomal trafficking and vesicle maturation [35], which may buffer or mask RNF13-driven effects. While Rab7-positive late endosomes were unaffected (Fig. 5K), cells expressing RNF13 L244A displayed a significant increase in peripheral Lamp1-positive lysosomes (Fig. 5L). Together, these findings identify Leu244 as a critical residue mediating the binding between RNF13 and Arl8B, suggesting that this interaction facilitates RNF13 targeting to lysosomes and contributes to the regulation of early endosome and lysosome positioning. Although not essential for RNF13’s localization to late compartments, disruption of this interaction, either through mutation or Arl8B depletion, reveals a context-dependent role for RNF13 in modulating the spatial organization of endolysosomal compartments. Furthermore, the altered lysosomal distribution associated with RNF13 L244A expression implies that the RNF13-Arl8B interaction contributes to lysosome positioning, potentially by stabilizing RNF13 association with Arl8B- positive lysosomes and limiting their peripheral redistribution under basal conditions.

### Arl8B interaction variants affect lysosomal properties and EGFR trafficking without altering the integrity of late endosomes

Thus, so far, this study has demonstrated that Arl8B variants exhibit altered subcellular localization and that Arl8B E22D and F55A variants specifically affect the positioning of early endosomes. We examined whether Arl8B variants affect the abundance of endolysosomal proteins. The presence of Arl8B WT, E22D, and F55A increased Lamp1 protein levels (Fig. 6A-B). The presence of Arl8B variants reduced EEA1 levels, whereas Rab7 levels remained constant across all groups (Fig. 6A-B). Next, to assess the functional relevance of these changes, the impact of Arl8B E22D and F55A variants on EGFR endocytosis and trafficking to lysosomes was measured by using a fluorescently labelled EGF complex. While there was no change in EGFR protein abundance (Fig. 6A-B), exogenous Arl8B WT and variants expressed in HeLa cells accelerated the initial EGF uptake compared to control cells (Fig. 6C). However, only Arl8B WT-expressing cells maintained higher residual EGF fluorescence at later timepoints (Fig. 6C). In contrast, cells expressing E22D and F55A variants showed enhanced internalization similar to vector-transfected cells (Fig. 6C). These results show that Arl8B E22D and F55A variants do not alter EGF internalization and degradation. Subsequently, we investigated whether Arl8B’s binding-defective variants impact the number and size of endolysosomal vesicles. The size of Rab7-positive late endosomes remained unchanged across conditions (Fig. 6D). In contrast, the presence of Arl8B E22D and F55A variants resulted in a decrease in lysosome size compared to Arl8B WT (Fig. 6E). A reduction in Rab7- positive vesicle numbers was also measured (Fig. 6F), whereas the number of Lamp1 puncta was unaltered (Fig. 6G). No changes were observed in the number or size of EEA1-positive early endosomes (data not shown). Altogether, these results suggest that Arl8B interaction variants alter lysosomal properties, including size and protein content, and modulate early stages of EGFR trafficking. These changes occur without affecting the morphology of late endosomes, providing further insight into how Arl8B interaction specificity regulates lysosomal organization and trafficking dynamics.

**Figure 6:**
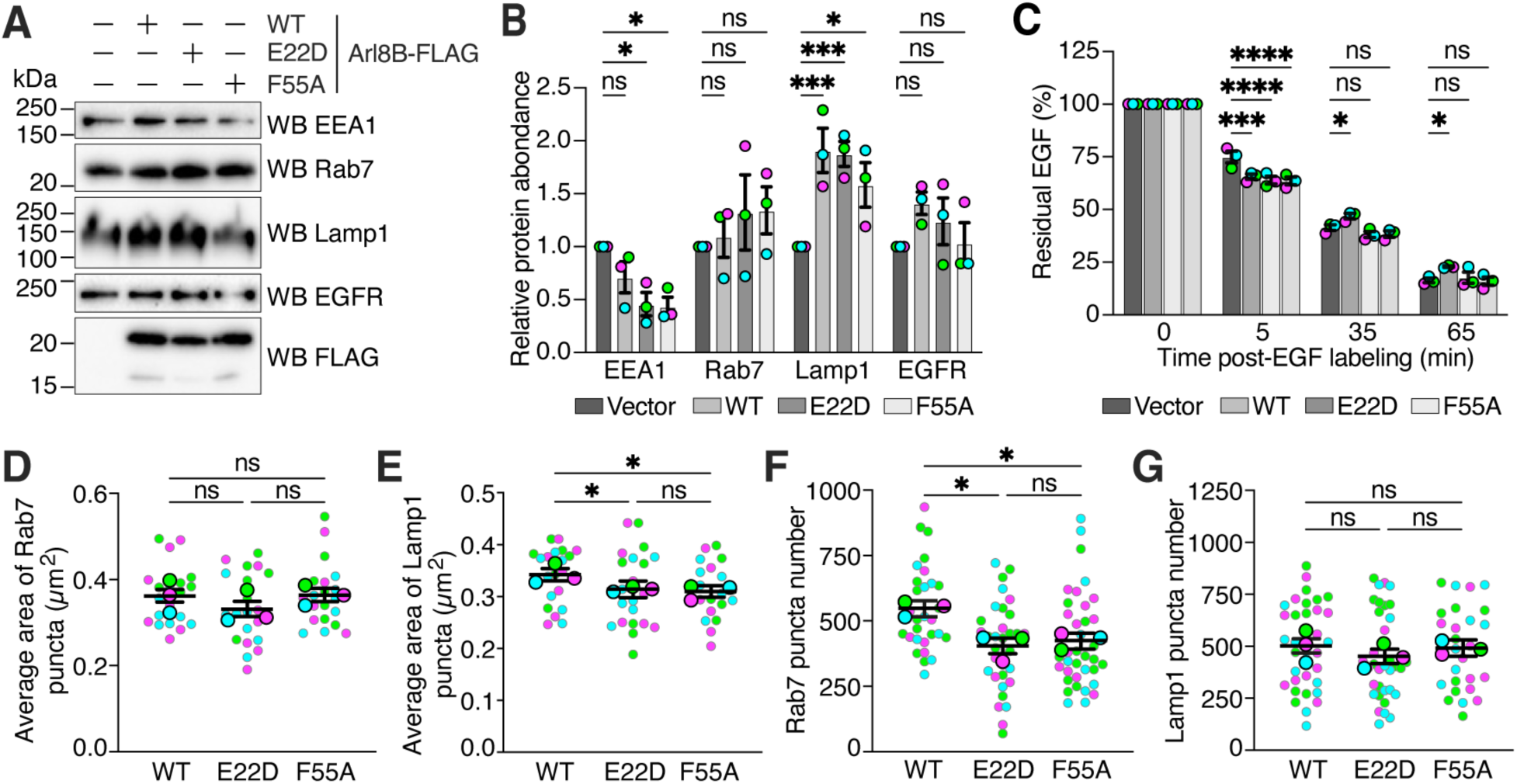
Arl8B binding-defective variants affect endolysosomal protein abundance and EGFR trafficking. (**A**) Representative (N=3) immunoblots show the effect of the ectopic expression of Arl8B binding-defective variants in HEK293T/17 cells on endolysosomal protein abundance. (**B**) The strip plot represents the normalized relative protein levels obtained after expression of Arl8B-FLAG (WT, E22D and F55A) in each independent experiment analyzed. (**C**) The strip plot represents the percentage of the average number of remaining EGF puncta per cell (n=35–45 cells per condition) at indicated time points. (**D-G**) The jittered individual value plots illustrate the effects of Arl8B binding-defective variants on the (**D-E**) average vesicle area and (**F-G**) the normalized vesicle counts of indicated markers per cell. Each color represents an independent experiment (N=3), with 10 cells analyzed per experiment (n=30). Larger circles indicate the experimental mean while smaller circles represent individual cell measurements. All results are expressed as mean ± SEM from N=3. Statistical significance was assessed using (**B, D-G**) one-way or (**C**) two-way RM-ANOVA with multiple comparisons test. ns: not significant; * *p* < 0.05; *** *p* < 0.001; **** *p* < 0.0001.

### RNF13-Arl8B interaction modulates EGF dynamics without altering the endolysosomal compartment

To further investigate the role of the RNF13-Arl8B interaction in endolysosomal system dynamics, we analyzed the effects of RNF13 WT and the interaction-defective RNF13 L244A mutant on several parameters related to endosomal and lysosomal compartments. In contrast to the results obtained with Arl8B variants, no significant differences in the number of EEA1, Rab7 or Lamp1 puncta were observed between vector control, RNF13 WT and RNF13 L244A- expressing cells (data not shown). Additionally, after assessing potential changes in protein levels or compartment content, the results from analyzing the fluorescence intensity of these markers did not show any significant changes across groups (data not shown). Thus, these results indicate that disrupting the interaction between RNF13 and Arl8B does not affect the steady-state abundance or spatial organization of key endolysosomal markers. Nonetheless, we evaluated whether the RNF13 L244A variant impacted the abundance of endolysosomal proteins in HEK293T/17 cells. Consistent with the imaging data, protein levels of EEA1, Rab7, and Lamp1 were comparable between conditions (Fig. 7A-B). However, RNF13 L244A expression resulted in a modest but statistically significant increase in EGFR protein abundance relative to vector control (Fig. 7A-B), suggesting altered receptor trafficking or degradation. Then, we assessed whether RNF13 L244A influenced the trafficking and degradation kinetics of fluorescently labelled EGF. By measuring fluorescence at an early time point (t = 5 minutes) following ligand internalization, the cells expressing RNF13 (WT or L244A) showed a significant increase in residual EGF compared to control cells (Fig. 7C), suggesting that RNF13 enhances early endocytic uptake. However, at 65 minutes post-labelling, RNF13 WT-expressing cells exhibited a significantly lower EGF signal compared to RNF13 L244A-expressing cells (Fig. 7C), indicating that the L244A substitution in RNF13 impairs EGF degradation. Taken together, these findings suggest that RNF13 promotes early endocytic uptake of EGFR and that its interaction with Arl8B may support efficient cargo trafficking to lysosomes, without altering endolysosomal compartment composition.

**Figure 7:**
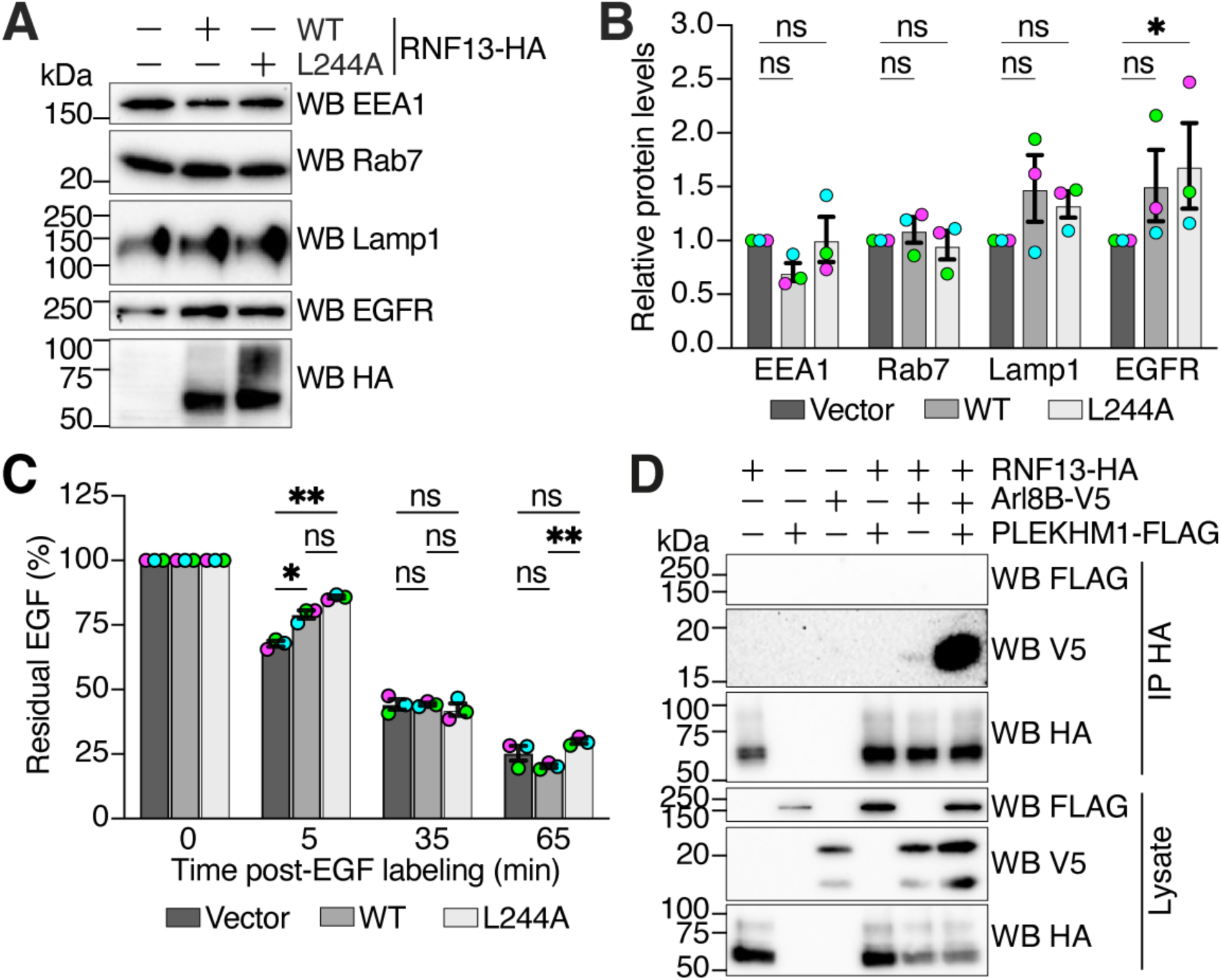
RNF13 binding-defective variant affects EGFR trafficking. (**A**) Representative (N=3) immunoblots show the effect of ectopic expression of RNF13 (WT and L244A) in HEK293T/17 cells on endolysosomal protein abundance. (**B**) The strip plot represents the normalized relative protein levels obtained after expression of RNF13-HA (WT and L244A) in each independent experiment analyzed. (**C**) The strip plot represents the percentage of the average number of remaining EGF puncta per cell (n=35–45 cells per condition) at the indicated time points. Each color represents an independent experiment (N=3), and large circles indicate the experimental mean. (**D**) Representative (N=4) immunoblots of co-immunoprecipitation assay showing that PLEKHM1-FLAG enhances the interaction between RNF13-HA and Arl8B-V5. Indicated plasmid constructs were co-expressed in HEK293T/17 cells, and HA immunoprecipitation was performed. All results are expressed as mean ± SEM from N=3. Statistical significance was assessed using (**B**) one-way or (**C**) two-way RM-ANOVA with multiple comparisons test. ns: not significant; * *p* < 0.05; ** *p* < 0.01.

Given the observed importance of the RNF13-Arl8B interaction for efficient EGFR trafficking, and the established role of Arl8B effectors in coordinating cargo delivery to lysosomes, we next explored whether another Arl8B binding partner might influence this interaction. Notably, both PLEKHM1 and RNF13 bind Arl8B through the identical critical residues (E22 and F55) [19], raising the question of whether these interactions are competitive or cooperative. To address this, a co-immunoprecipitation assay was performed using lysates expressing a combination of RNF13-HA, Arl8B-V5 and PLEKHM1-FLAG. Strikingly, the presence of PLEKHM1 increased the co-precipitation of Arl8B with RNF13 compared to the condition where only RNF13 and Arl8B were expressed (Fig. 7D). This suggests that PLEKHM1 enhances the RNF13-Arl8B interaction, rather than sequestering Arl8B from RNF13, highlighting a potential cooperative mechanism in lysosomal trafficking.

Together, these findings indicate that disruption of the RNF13-Arl8B complex through the RNF13 L244A variant does not impact the structural organization of endolysosomal compartments. However, this interaction functionally contributes to EGFR trafficking, as its loss results in delayed degradation of the internalized ligand. These results suggest that RNF13 Arl8B binding facilitate the efficient progression of internalized cargo toward lysosomal compartments. Moreover, PLEKHM1 may facilitate the formation of an RNF13-Arl8B complex, revealing an additional regulatory layer in the control of endolysosomal trafficking.

### RNF13 interacts with Arl8B in both GDP- and GTP-bound conformations

To further understand the nature of the RNF13-Arl8B interaction, we examined whether Arl8B’s activation state influences RNF13 binding. While established effectors such as PLEKHM1, SKIP and RUFY3 are known to bind specifically to the GTP-bound form of Arl8B, consistent with the classic mode of effector recognition by small GTPases [10, 12, 14, 19], it is unknown whether RNF13 engages Arl8B in a manner consistent with these effectors. To explore this, we used Alphafold3 to predict the structural interface of RNF13 in association with Arl8B in both GTP- and GDP-bound conformations. Remarkably, RNF13 was predicted to bind both forms, with stable interfaces observed in the Alphafold3 models (Fig. 8A-B), suggesting that RNF13 does not exhibit a strict preference for either Arl8B activity state. Although the Predicted Aligned Error (PAE) confidence levels of both predictions were slightly lower than the original prediction with AlphaFold 2, the interactions involve the same binding interface and a very similar set of residues. To validate these predictions, we performed co-immunoprecipitation assays using RNF13 and Arl8B activity variants, the constitutively active Q75L variant (GTP-locked) and the constitutively inactive T34N variant (GDP-locked). Consistent with the *in-silico* models, RNF13 was able to co-immunoprecipitate with both Arl8B variants (Fig. 8C), indicating that its interaction is not dependent on Arl8B’s nucleotide-bound state. These findings distinguish RNF13 from classical Arl8B effectors and support a model in which RNF13 may function independently of Arl8B’s activation cycle, potentially serving as a platform that modulates or stabilizes vesicular interactions regardless of Arl8B’s conformational state. To explore this possibility, we propose a conceptual model (Fig. 8D) in which RNF13 associates with Arl8B across its nucleotide cycle, potentially supporting distinct stages of lysosomal trafficking. This model raises the question of how RNF13 may modulate Arl8B-dependent pathways and whether its interaction influences effector recruitment, vesicle identity or trafficking efficiency. Rather than acting as a conventional effector that selectively binds the active form, RNF13 appears to interact with Arl8B independently of its activation state, possibly serving as a scaffold or stabilizer throughout its functional cycle.

**Figure 8:**
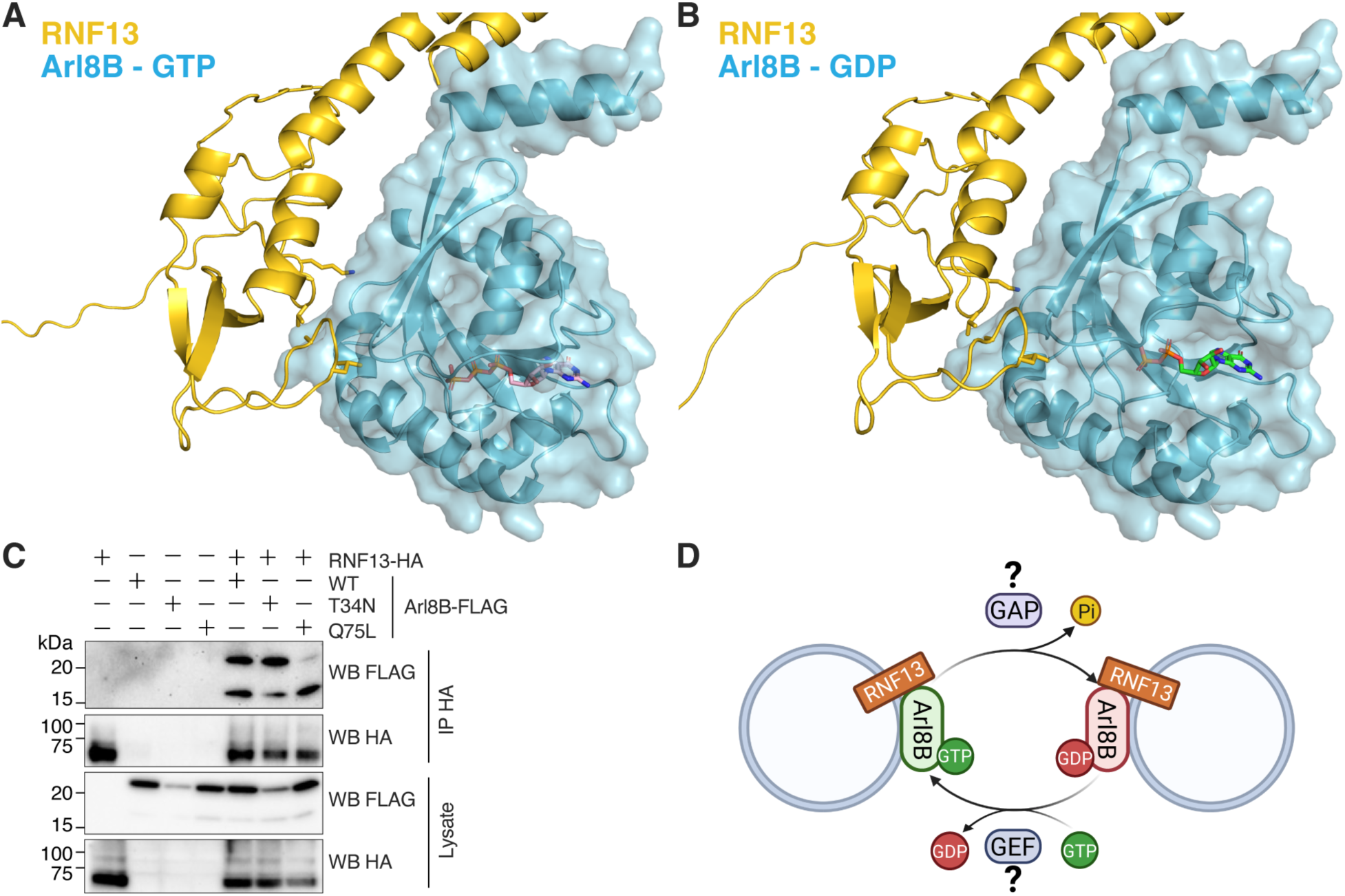
RNF13 interacts with Arl8B regardless of its activity state. (**A-B**) AlphaFold3-predicted structures of complexes between RNF13 (yellow) and Arl8B (blue) in its (**A**) GTP-bound and (**B**) GDP-bound conformations. RNF13 is shown in stick representation and Arl8B in surface representation. (**C**) Representative (N=3) immunoblots of co-immunoprecipitation from HEK293T/17 cells transiently transfected with HA-tagged RNF13 and FLAG-tagged Arl8B activity variants (constitutively active Arl8B Q75L (GTP-locked) and inactive Arl8B T34N (GDP-locked)) showing RNF13 associates with both forms. (**D**) Hypothetical model illustrating the nucleotide cycle of Arl8B and the binding of RNF13 in both GDP- and GTP-bound states. RNF13 may serve as a scaffold that associates with Arl8B regardless of its activation status, supporting vesicle positioning or trafficking across multiple stages of the endolysosomal pathway. Created in BioRender. Sénécal, A. (2025) https://BioRender.com/s4uu3ws.

## DISCUSSION

This study uncovers a novel role for RNF13 in modulating endolysosomal dynamics through its interaction with the small GTPase Arl8B. While RNF13 has been linked to the ubiquitin-mediated degradation of targets such as Lamp1 and SNAPIN [29, 30], our findings indicate a ubiquitin-independent role for RNF13 in the endolysosomal pathway, suggesting a broader regulatory mechanism in intracellular trafficking. Our study demonstrates that RNF13 associates with Arl8B through a defined binding interface involving RNF13 residue Leu244 and Arl8B residues Glu22 and Phe55. These two residues on Arl8B are also needed for the recruitment of other Arl8B effectors such as PLEKHM1 and SKIP [19], indicating a shared binding interface among functionally distinct partners. This specificity may serve as a regulatory checkpoint, where RNF13 influences the effector landscape of Arl8B, thereby affecting vesicular trafficking outcomes in a context-dependent manner. Our data support a non-enzymatic contribution of RNF13 to lysosomal organization and function, independently of its catalytic activity. Disruption of the RNF13-Arl8B interaction impairs EGFR degradation, despite normal initial internalization, without altering the overall abundance or number of endolysosomal compartments. This suggests that RNF13 specifically facilitates the late stages of cargo progression, particularly lysosomal delivery and degradation, rather than having a broad impact on endosomal structure or uptake efficiency [28, 29]. Interestingly, EGFR internalization was increased in both RNF13 WT- and L244A-expressing cells. However, only RNF13 WT facilitated efficient degradation, suggesting that the RNF13-Arl8B complex promotes timely progression from early uptake to lysosomal delivery, rather than acting as a brake on trafficking.

The spatial distribution of lysosomes is tightly linked to their functional state, with perinuclear lysosomes being more degradative than those located at the cell periphery [36, 37]. Consistent with this, we observed that disruption of the RNF13-Arl8B interaction, either by knocking down RNF13 or expressing RNF13-binding-defective mutants of Arl8B, resulted in the repositioning of lysosomes toward the cell periphery. This phenotype reflects those reported for defects in dynein-mediated retrograde transport [14], suggesting that RNF13 may indirectly affect motor complex recruitment or activity, possibly by modulating Arl8B’s interaction with its bidirectional trafficking partners such as RUFY3, PLEKHM2 or PLEKHM1 [5, 10, 12]. In addition, despite these changes in lysosome positioning, neither Arl8B variants nor RNF13 L244A significantly altered the number, intensity, or global organization of endolysosomal markers such as EEA1, Rab7, or Lamp1. This mechanism represents a regulatory process that enables cells to quickly adapt the positioning of lysosomes in response to environmental or signaling cues, all without the need for extensive remodeling of the endolysosomal system. Interestingly, contrary to previous assumptions, the Arl8B interaction-deficient variants (E22D and F55A) exhibited increased colocalization with early endosomes, yet they did not alter the overall number or size of the early endosomal marker EEA1. This finding aligns with prior research by Rawat et al. (2023), which indicated that Arl8B primarily regulates lysosomal and late endosomal functions. It only colocalizes with a subset of early endosomes through effectors such as RUFY1 [38]. importantly, RUFY1 interacts with Arl8B on non-acidic compartment that are positive for Rab14 and EEA1. However, the depletion of Arl8B does not disrupt the overall architecture of early endosomes or the distribution of EEA1. These results indicate that Arl8B-binding-defective mutants affect lysosomal and EGFR trafficking without compromising the structural integrity of early endosomes. Instead, this reinforces the notion that Arl8B’s role in early endosomal compartments is context-specific, likely mediated by selective effector engagement with effectors like RUFY1, rather than providing broad structural control over early endosome identity.

Our study reveals a potential cooperative interaction between RNF13 and PLEKHM1 at the Arl8B interface. Co-expression of PLEKHM1 enhances the association of RNF13 with Arl8B, proposing a model where these effectors may bind sequentially or support each other’s interaction with Arl8B, depending on the cellular context. Since PLEKHM1 is crucial for HOPS complex assembly and lysosome-endosome fusion [13], it is possible that RNF13 plays a role in activating Arl8B or facilitates a more efficient transfer to downstream effectors, such as PLEKHM1. Alternatively, RNF13 may serve as a molecular scaffold, positioning Arl8B alongside other tethering factors or restricting access to competing effectors, thus influencing the sequence of recruitment events. This hierarchical assembly mechanism, in which upstream effectors influence the availability of small GTPases for downstream interactions, is crucial for Arl8B and its regulation by RNF13. A notable example of this is the transition from Rab7 to Arl8B during the maturation of late endosome to lysosome. Recent study indicates a temporal and spatial separation between these two GTPases where Rab7 mainly handles late endosomal functions, while Arl8B is responsible for lysosomal positioning and fusion [34]. This transition is likely regulated by effector exchange and compartment maturation, where Rab7 effectors like RILP, coordinate degradative transport, whereas Arl8B effectors, such as PLEKHM1 and SKIP facilitate lysosomal movement and fusion [9, 10, 39]. The sequential recruitment of these small GTPases exemplifies the potential role of RNF13 in modulating Arl8B function during this maturation process, suggesting that RNF13 orchestrates the transition between different states of effector complexes. The enhanced interaction between RNF13 and Arl8B, driven by co-expression of PLEKHM1, further indicates that the RNF13- Arl8B complex adapts to cargo trafficking needs.

A particularly striking finding was the increased peripheral of Rab7 localization observed upon RNF13 depletion combined with Arl8B overexpression, contradicting the model where Arl8B displaces Rab7 via the SKIP–HOPS–TBC1D15 complex, leading to Rab7 inactivation and removal from lysosomal membranes [9, 10]. While the mechanisms underlying this altered Rab7 distribution remain unclear, it may result from the interplay between RNF13 and RNF167, which has been shown to heterodimerize and influence lysosomal trafficking [40]. Specifically, RNF167 regulates endosomal and lysosomal positioning by controlling the activities of Rab7 and Arl8B [22, 41]. A recent study found that RNF13 positively regulates RNF167’s trafficking to Lamp1-positive lysosomes, and its depletion reduces RNF167’s lysosomal association [40]. The loss of RNF13 may promote RNF167 homodimer formation or enhance its activity, leading Rab7 retention at peripheral endosomes despite Arl8B overexpression. This model suggests that RNF167 counters Rab7 displacement by Arl8B, either by stabilizing active Rab7 on membranes or by delaying its inactivation. The combined effects of Arl8B-induced vesicle dispersion and enhanced RNF167-driven Rab7 retention result in increased Rab7 presence on peripheral late endosomes, explaining the unexpected Rab7 localization pattern. This mechanism underscores the importance of the RNF13–RNF167 axis in spatially regulating the endolysosomal system, suggesting a role in modulating the balance between Rab7 and Arl8B compartmentalization.

Our data indicate that RNF13 is important not only for vesicle positioning but also for guiding the cargo fate within the endolysosomal network, aligning with studies that show spatial regulators of lysosomes influence receptor destiny [10, 38, 42]. Cells expressing RNF13 variants that cannot interact with Arl8B showed defects in EGFR trafficking and degradation, underscoring the importance of this interaction in the progression of cargo from initial uptake to degradation-competent lysosomes. This is particularly relevant given the role of lysosomal degradation in regulating signaling receptor turnover. Additionally, disrupting the RNF13-Arl8B interaction through the expression of RNF13-binding-defective Arl8B variants altered lysosomal positioning and impaired EGFR degradation, thereby affecting the abundance of endosomal compartments. These results suggest that RNF13’s role is not in compartment biogenesis, but rather in enhancing cargo processing and fusion events. Its interaction with Arl8B likely functions as a checkpoint, coordinating vesicle maturation and spatial distribution with the timely degradation of internalized receptors. Therefore, RNF13 may act as a modulator of fusion competency or as an effector during the endolysosomal maturation process. Consequently, RNF13 emerges as a key regulator of lysosomal homeostasis, especially under conditions that necessitate coordinated trafficking responses, such as increased receptor turnover or cellular stress.

In conclusion, this study identifies RNF13 as a novel modulator of endolysosomal trafficking, acting not through its enzymatic activity, but by scaffolding specific molecular interactions. By characterizing the interaction of RNF13 with Arl8B under basal conditions and examining the downstream effects on lysosome positioning and cargo trafficking, we reveal a new layer of complexity in vesicular trafficking regulation. RNF13 thus emerges as a key regulator of endolysosomal dynamics and a potential contributor to cellular homeostasis. Future research should focus on defining the dynamic regulation of the RNF13-Arl8B axis, its specific roles in different cellular contexts, and its interactions with other trafficking machinery to achieve a fully comprehensive understanding of RNF13’s biological functions.

## MATERIALS AND METHODS

### Antibodies

The following antibodies were used: rabbit anti-HA 1.1 (WB and IF: 1:1000, BioLegend, San Diego, CA, USA, Cat# 902302), mouse anti-HA 1.1 (IF: 1:1000, BioLegend, San Diego, CA, USA, Cat# 901503), Rabbit anti-RNF13 (WB: 1:500, Cusabio Technology LLC, Cat# CSB-PA019831DSR1HU), Mouse anti-Rab7 (IF: 1:750, New England Biolabs Ltd., Cat# 957468), rabbit anti-Lamp1 (IF: 1:500, New England Biolabs Ltd., Cat# 9091, RRID:AB_2687579), rabbit anti-EEA1 (IF: 1:500, New England Biolabs Ltd., Cat# 3288), rabbit anti-EGFR (IF: 1:1000, New England Biolabs Ltd., Cat# 4267), mouse anti-FLAG (WB and IF: 1:1000, SigmaMillipore Canada, Mississauga, ON, Canada, Cat# F1804), rabbit anti-Flag (WB and IF: 1:1000, New England Biolabs Ltd., Cat# 14793), mouse anti-P4D1 (1:1000, Santa Cruz Biotechnology, Cat# sc-8017), AlexaFluor Plus 488-conjugated goat anti-rabbit (1:1000, Thermo Fisher Scientific, Cat# A32731), AlexaFluor Plus 488-conjugated goat anti-mouse (1:1000, Thermo Fisher Scientific, Cat# A32723), AlexaFluor Plus 647-conjugated goat anti-rabbit (1:1000, Thermo Fisher Scientific, Cat# A32733), AlexaFluor Plus 647-conjugated goat anti-mouse (1:1000, Thermo Fisher Scientific, Cat# A32728), horseradish peroxidase (HRP)- conjugated goat anti-mouse (1:10000, New England Biolabs Ltd., Cat# 7076, RRID:AB_330924) and horseradish peroxidase (HRP)-conjugated goat anti-rabbit (1:10000, New England Biolabs Ltd., Cat# 7074, RRID:AB_2099233).

### Molecular Biology

pCAGGs-IRES-mCherry plasmid encoding for full-length human RNF13 transcript variant 1 (accession NM_007282.4) containing an optimized Kozak sequence and a C-terminus HA tag was obtained as previously described [28]. pcDNA3.1+/C-(K)DYK plasmid encoding the full-length human Arl8B transcript variant 1 (accession NM_018184.3) was purchased from GenScript (GenScript, CloneID #OHu02123) and served as template for site-directed mutagenesis of E22D, F55A, T34N and Q75L using SSB-Fusion polymerase or subsequent subcloning in pmCherry-N vectors, using standard restriction enzymes strategies followed by ligation. pcDNA3.1(+) plasmid encoding full-length human PLEKHM1 (accession NM_183034) containing a C-terminus FLAG tag was a gift from Paul Odgren (Addgene plasmid # 73593; http://n2t.net/addgene:73593; RRID: Addgene_73593) [43]. All plasmids were confirmed by Sanger sequencing (Genome Quebec, Montreal, QC, Canada).

### Cell Culture and Transfection

HeLa (from Dr. Diana Alison Averill, Montréal) and HEK293T/17 (American Type Culture Collection (ATCC), Gaithersburg, MD, USA, Cat# CRL- 11268) cells were cultured in Dulbecco’s Modified Eagle Medium (DMEM, ThermoFisher Scientific, Cat# 11995-065) supplemented with 10% fetal bovine serum (FBS, Corning, Cat# 35-077-CU) in an incubator at 37°C with a humidified atmosphere containing 5% CO_2_. For the immunoprecipitation assays, HEK293T/17 cells were transfected in poly-D-lysine treated six-well plate using LipofectAMINE 2000 as described previously [44]. Cells were maintained for 24 hours at 37°C with 5% CO_2_ before being lysed. For immunostaining assay, 1.25 x 10^5^ HeLa cells were seeded 24 hours before transfection in a 24-well plate containing a 12 mm round glass coverslip #1.5 (UltiDent Scientific Inc., St-Laurent, QC, Canada, Cat# 170-C12MM). Cells were transfected with a calcium phosphate-DNA precipitate as described previously [28] and then maintained for 20-24 hours at 37°C with 5% CO_2_ before being fixed and processed for immunostaining.

### Arl8B and RNF13 Knockdown

TriFECTa Kit Dicer-substate siRNAs duplexes against Homo sapiens ADP-ribosylation factor-like protein 8B (Arl8B, NM_018184) and RING-finger protein 13 (RNF13, NM_007282) were purchased from Integrated DNA Technology (IDT, Coralville, IA). The non-targeting DsiRNA, DS NC1 was used as a universal negative control (Scrambled Negative Control DsiRNA, IDT, Coralville, IA, Cat# 51-01-19-08). Precisely, DsiRNAs against Arl8B sequences (#1: 5’- rGrGrArGrUrCrArArUrGrCrUrArUrUrGrUrU-3’ and 5’-rUrCrArUrGrUrArArArCrArArUrArGrCrArU-3’, and #2: 5’-rArCrArCrUrGrArCrUrUrGrArUrArCrArGrC-3’ and 5’- rArCrUrUrCrArUrGrCrUrGrUrArUrCrArArG-3’) and against RNF13 sequences (#1: 5’rGrGrUrGrArArUrCrArUrCrArGrCrUrArA-3’ and 5’- rUrCrArGrArGrArGrArArUrUrArGrCrUrGrArU-3’, and #3: 5’-rCrGrArCrArUrUrGrArGrGrUrArC- 3’ and 5’-rGrUrArGrUrArCrCrUrCrArArUrG-3’) were used. Briefly, 1.25x10^5^ Hela cells were plated 2 hours before transfection in a 24-plate well containing a 12mm round glass #1.5 coverslip. Then, the cells were transfected using a mixture of 0.5uL LipofectAMINE 2000 transfection reagent (Thermo Fisher Scientific, cat# 11668-019, Lot# 2531744) diluted in 50uL Extreme MEM (Wisent, Saint-Jean-Baptiste, QC, Canada, Cat# 390-005-CL, Lot# 39005033) which was added to a total of 10 nM DsiRNAs in 50uL ExtremeMEM. The mixture was incubated for 20 minutes at room temperature and added to the cells. Finally, 48 hours after transfection with DsiRNAs, cells were transfected for 24h with the plasmid constructs encoding RNF13-HA or Arl8B-FLAG using a calcium phosphate-DNA precipitate.

### Immunostaining

Transfected HeLa cells were washed three times in phosphate-buffered saline (PBS) before being fixed for 15 mins with 4% paraformaldehyde/4% sucrose in PBS at room temperature. Fixed cells were washed three times and permeabilized with 0.25% Triton X-100 in PBS (PBS-T) for 15 mins at room temperature. Non-specific sites were blocked using a blocking solution composed of 10% normal goat serum in PBS for 1 hour. Staining was performed immediately after incubating with primary antibodies diluted in 3% NGS/PBS for 1 hour. Coverslips were washed three times with PBS, followed by incubation with the appropriate conjugated secondary antibodies diluted in 3% NGS/PBS for 1 hour. After being washed in PBS, coverslips were mounted using ProLong Diamond Antifade (Thermo Fisher Scientific, Cat# P36961) and dried overnight at room temperature. Image acquisitions were made the next day.

### Fluorescence Microscopy Image Acquisition

Image acquisition was performed using an inverted epi-fluorescence microscope Olympus IX83 equipped with a U plan S-Apo 60x / 1.35 numerical aperture oil objective (Olympus), the X-Cite Xylis 365 LED-based illumination source (Excelitas Technologies Corp., Waltham, MA, USA) and a Zyla 4.2 Plus sCMOS camera (Andor, Oxford, UK). Olympus CellSens Dimension software version 2.2 (Olympus, Toronto, ON, Canada, RRID: SCR_014551) controlled the system during image acquisition. For fixed cells, the resolution was set at 2048 x 2048 pixels, and z-stack images were taken at 0.27 µm intervals for 6 to 11 optical slices. Lamp intensity and exposure time were set to obtain the highest signal possible without reaching saturation. Before analysis, images were deconvoluted using the Olympus 3D Deconvolution feature in the Olympus CellSens Dimension software. For live-cell imaging, the resolution was set at 1024 x 1024 pixels, with constant lamp intensity and exposure time for each independent experiment. The Olympus Z Drift Compensator 2 hardware module (IX3-ZDC2) was used to enable continuous autofocus during the acquisition of a single z-slice over time. Fluorescence signals were acquired every 3 s for 100 cycles (a total of 5 min), starting by mCherry followed by EGFP.

### Live-cell imaging

Transfected HeLa cells were washed once with PBS and incubated in FluoroBrite DMEM (Thermo Fisher Scientific, Cat# A1896701) supplemented with 10% FBS and 2mM L-Glutamine 20 minutes before the acquisition. Coverslips were mounted in an Attofluor Cell Chamber (Thermo Fisher Scientific, Cat# A7816) with 1mL of supplemented FluoroBrite DMEM and placed in a warm humidified stage-top incubator (Okolab UNO-T1) coupled with an objective heating collar (Okolab), keeping the live cell sample at 37°C with 5% CO_2_ during the acquisition.

### Co-immunoprecipitation Assays

Transfected HEK293T/17 cells were washed twice with PBS and lysed in 1 mL ice-cold lysis buffer (20mM Tris-HCl pH 7.5, 150 mM NaCl, 1 % Triton X-100, 0.5 % sodium deoxycholate and 1X protease inhibitor cocktail EDTA-free (Bimake.com, Houston, TX, USA, Cat# B14001)). Cell lysates were collected, agitated at 15 rpm for 20 mins at 4°C and centrifuged at 21,000 x g for 15 mins. For co-immunoprecipitation with RNF13-HA and Arl8B-Flag, 700 µL of total lysate was agitated at 15 rpm for 2h at room temperature with 10 µL of anti-HA magnetic beads (Bimake.com, Houston, TX, USA, Cat# B26201) pre-equilibrated in PBS-T or for 2h at 4°C with 5uL of anti-FLAG^®^ M2 affinity gel (Sigma-Aldrich, Missouri, USA, Cat# A2220) pre-equilibrated in ice-cold lysis buffer. After co-immunoprecipitation with anti-HA beads, protein complexes were washed 3x 5 mins with PBS- T, and proteins were eluted in 1X Laemmli buffer and boiled for 5 mins at 95°C before loading on SDS-PAGE. Besides, protein complexes co-immunoprecipitated with anti-FLAG beads were washed three times with 1mL of ice-cold lysis buffer before being eluted by competitive elution using 6 µg of 3X FLAG^®^ Peptide (Sigma-Aldrich, Missouri, USA, Cat# F4799) in water for 30mins at 37°C and 1X Laemmli buffer was added prior to SDS-PAGE loading.

### SDS-PAGE and Western Blot

Protein samples were separated on a 1.5 mm SDS-PAGE gel containing 0.5% of 2,2,2-trichloroethanol (TCE) and transferred to a 0.45 µm PVDF membrane using Bio-Rad Trans-Blot Turbo system (10 mins, constant 2.5 A, 25V). Proteins on both gels and membranes were visualized using the *stain-free* mode of the ChemiDoc MP imaging System (Bio-Rad) and Bio-Rad Image Lab software. Non-specific sites were blocked using 5% skim milk in TBS-T (20 mM Tris-HCl pH 7.5, 140 mM NaCl, 0.3 % Tween-20) for 1h at room temperature. Membranes were incubated overnight at 4°C with primary antibodies diluted in TBS-T containing 0.05 % NaN_3_. Membranes were washed three times with TBS-T before being incubated for 1h at room temperature with horseradish peroxidase-coupled secondary antibodies diluted in TBS-T. Immune complexes were revealed using the Clarity Western ECL substrate (BioRad, Cat# 1705060). The ChemiDoc MP imaging system was used to acquire the chemiluminescent signal.

### Analysis of endolysosomal compartments abundance

To assess the impact of the loss of interaction on the abundance of the various intracellular compartments, HEK293T were transfected with plasmid constructs encoding for Arl8B-FLAG (WT, E22D or F55A) or empty vector for 24h. Cells were washed in PBS and lysed with ice-cold RIPA lysis buffer (20mM Tris-HCl pH 7.5, 150 mM NaCl, 1 % Triton X-100, 5mM Ethylenediaminetetraacetic acid (EDTA), 0.1% Sodium Dodecyl Sulfate (SDS), 0.5 % sodium deoxycholate and 1X protease inhibitor cocktail EDTA-free (Bimake.com, Houston, TX, USA, Cat# B14001)). Cell lysates were obtained as described previously, diluted in 1X Laemmli buffer, and loaded onto SDS-PAGE for Western blotting analysis. ImageJ software was used to quantify the relative protein abundance of the markers through densitometric analysis of immunoblots, which were normalized to the stain-free signal and then to the control condition.

### EGF uptake and internalization assay

For EGF internalization and trafficking assay, transfected HeLa cells were washed with PBS and starved for 2h at 37°C in serum-free DMEM supplemented with 0.2% bovine serum albumin. Then, cells were placed on ice, the medium was replaced by 0.5 µg/mL AlexaFluor™647 EGF complex (Thermo Fisher Scientific, Cat# E35351) in DMEM+0.2% BSA medium and the cells were incubated for 10 minutes, always on ice. After the incubation period, cells were kept on ice and washed with ice-cold PBS to remove traces of labeling solution while inhibiting endocytosis. Following washes, cells were incubated in fresh culture medium (DMEM containing 10% FBS) at 37°C to induce EGF internalization and cells were incubated for a period corresponding to each time point. Immediately after each timepoint, cells were washed in PBS and fixed with ice-cold 4% PFA/4% sucrose before performing immunostaining and image acquisition as previously described.

### Image Analysis and Statistical Analysis

Colocalization assays were analyzed on a single slice of the z-stack for each image after separating the channels. The threshold was adjusted to obtain a minimal background and Mander’s overlap coefficient was measured using the JaCoP plugin in Fiji. The distribution, number of vesicles, Pearson’s overlap correlation coefficient and the per cell average individual vesicle size and intensity analysis were made with CellProfiler 4.2.6 on a single Z-plane of the images. A CellProfiler pipeline was created to quantify the previously mentioned parameters, and all cell images were analyzed with it.

Briefly, images were converted to grayscale, and channels were split using the “ColorToGray” module. Nuclei and proteins of interest were identified using the IdentifyPrimaryObjects with a Global Otsu thresholding method. Cell boundaries were manually identified. Primary object measurements were performed, and the values obtained for each object were linked to the corresponding cells using the “RelateObjects” module. The distribution was analyzed by measuring the intensity distribution, specifically the fraction at a distance (FracAtD), from the nucleus to the cell periphery of the marker, representing the fraction of total stain in an object at a given radius. Briefly, each cell was divided into 10 bins extending from the nucleus to the edge of the cell (with bin widths normalized per cell), and the total intensity within each fraction was measured. Ultimately, the “ExportToSpreadsheet” module exported all the measurement values to an Excel spreadsheet. For the distribution values, the perinuclear region corresponds to the sum of bins 1 and 2, while the peripheral region corresponds to the sum of bins 9 and 10. At least three independent, non-blinded experiments were performed for all assays. Datasets of image analysis were exported to a CSV file and collected in Microsoft Excel before being subjected to statistical analysis in GraphPad Prism 10. Parametric tests were used because the data followed a Gaussian distribution. Statistical difference was determined using either repeated-measures one-way ANOVA with Dunnett’s or Turkey’s multiple comparisons test when there is one grouping variable or two-way ANOVA with Dunnett’s multiple comparisons test when there are two grouping variables.

### Predicted Structure

The structures of the complex formed by RNF13 (Uniprot accession O43567) and Arl8B (Uniprot accession Q9NVJ2) was modeled using ColabFold/AlphaFold2 using computational resources of Calcul Canada [45]. The structures of the complex formed by RNF13 (Uniprot accession O43567) and Arl8B (Uniprot accession Q9NVJ2) in complex with either GTP or GDP was modeled using AlphaFold3 [46]. The RNF13–Arl8B complexes were visualized using the PyMOL Molecular Graphics System, Version 3.1.3 Schrödinger, LLC. The models are available in ModelArchive (www.modelarchive.org) with the accession codes ma- cs3ca (Figure 1A and 4A), ma-hp6d3 (Figure 8A) and ma-tlkyz (Figure 8B) [47].

## ACKNOWLEDGEMENTS

This work fulfills part of the requirements for AMS’s PhD thesis at the *Université du Québec à Montréal*. Funding was provided by the *Centre d’Excellence en Recherche sur les Maladies Orphelines—Fondation Courtois* (CERMO-FC) for New Collaborative Research Initiatives (to MPL and LC), and through graduate scholarships for AMS and VCC. This research was partially funded by a Discovery Grant from the Natural Sciences and Engineering Research Council of Canada (RGPIN-2017-05392). The *Regroupement québécois de recherche sur la fonction, l’ingénierie et les applications des protéines* (PROTEO, DOI: https://doi.org/10.69777/341121) awarded PhD graduate scholarships to AMS. The FRQNT provided PhD graduate scholarships to VCC (DOI: https://doi.org/10.69777/326578) and AYB (DOI: https://doi.org/10.69777/317082). Figure 8D was created with BioRender.com under a publication license (https://BioRender.com/s4uu3ws). The authors declare that the funding sources had no role in study design, data collection, analysis, or interpretation.

## ABBREVIATIONS

Arl8B: ADP-ribosylation factor-like protein 8B
RNF13: RING finger protein 13
PA-TM-RING: Protease-Associated, Transmembrane, RING Finger
EEA1: Early Endosome Antigen 1
EGF: Epidermal Growth Factor
EGFR: Epidermal Growth Factor Receptor
Lamp1: Lysosomal Associated Membrane Protein 1
PLEKHM: Pleckstrin Homology Domain Containing Family M Member
PN: Perinuclear
PP: Peripheral
Rab7: Ras-related protein Rab-7a
RM: Repeated measures
RUFY: RUN and FYVE Domain Containing
SEM: Standard Error of the Mean

## REFERENCES

1. Klumperman, J. & Raposo, G. (2014) The complex ultrastructure of the endolysosomal system, Cold Spring Harb Perspect Biol. 6, a016857.

2. Huotari, J. & Helenius, A. (2011) Endosome maturation, Embo j. 30, 3481–500.

3. Singh, J., Elhabashy, H., Muthukottiappan, P., Stepath, M., Eisenacher, M., Kohlbacher, O., Gieselmann, V. & Winter, D. (2022) Cross-linking of the endolysosomal system reveals potential flotillin structures and cargo, Nature Communications. 13, 6212.

4. Winckler, B., Faundez, V., Maday, S., Cai, Q., Guimas Almeida, C. & Zhang, H. (2018) The Endolysosomal System and Proteostasis: From Development to Degeneration, J Neurosci. 38, 9364–9374.

5. Kumar, G., Chawla, P., Dhiman, N., Chadha, S., Sharma, S., Sethi, K., Sharma, M. & Tuli, A. (2022) RUFY3 links Arl8b and JIP4-Dynein complex to regulate lysosome size and positioning, Nat Commun. 13, 1540.

6. Filipek, P. A., de Araujo, M. E. G., Vogel, G. F., De Smet, C. H., Eberharter, D., Rebsamen, M., Rudashevskaya, E. L., Kremser, L., Yordanov, T., Tschaikner, P., Fürnrohr, B. G., Lechner, S., Dunzendorfer-Matt, T., Scheffzek, K., Bennett, K. L., Superti-Furga, G., Lindner, H. H., Stasyk, T. & Huber, L. A. (2017) LAMTOR/Ragulator is a negative regulator of Arl8b- and BORC-dependent late endosomal positioning, J Cell Biol. 216, 4199–4215.

7. Jean, S. & Kiger, A. A. (2012) Coordination between RAB GTPase and phosphoinositide regulation and functions, Nat Rev Mol Cell Biol. 13, 463–70.

8. Bonifacino, J. S. & Neefjes, J. (2017) Moving and positioning the endolysosomal system, Curr Opin Cell Biol. 47, 1–8.

9. Khatter, D., Raina, V. B., Dwivedi, D., Sindhwani, A., Bahl, S. & Sharma, M. (2015) The small GTPase Arl8b regulates assembly of the mammalian HOPS complex on lysosomes, J Cell Sci. 128, 1746–61.

10. Marwaha, R., Arya, S. B., Jagga, D., Kaur, H., Tuli, A. & Sharma, M. (2017) The Rab7 effector PLEKHM1 binds Arl8b to promote cargo traffic to lysosomes, J Cell Biol. 216, 1051–1070.

11. Kumar, R., Khan, M., Francis, V., Aguila, A., Kulasekaran, G., Banks, E. & McPherson, P. S. (2024) DENND6A links Arl8b to a Rab34/RILP/dynein complex, regulating lysosomal positioning and autophagy, Nature Communications. 15, 919.

12. Rosa-Ferreira, C. & Munro, S. (2011) Arl8 and SKIP act together to link lysosomes to kinesin-1, Dev Cell. 21, 1171–8.

13. McEwan, D. G., Popovic, D., Gubas, A., Terawaki, S., Suzuki, H., Stadel, D., Coxon, F. P., Miranda de Stegmann, D., Bhogaraju, S., Maddi, K., Kirchof, A., Gatti, E., Helfrich, M. H., Wakatsuki, S., Behrends, C., Pierre, P. & Dikic, I. (2015) PLEKHM1 regulates autophagosome-lysosome fusion through HOPS complex and LC3/GABARAP proteins, Mol Cell. 57, 39–54.

14. Keren-Kaplan, T., Sarić, A., Ghosh, S., Williamson, C. D., Jia, R., Li, Y. & Bonifacino, J. S. (2022) RUFY3 and RUFY4 are ARL8 effectors that promote coupling of endolysosomes to dynein-dynactin, Nature Communications. 13, 1506.

15. Cabukusta, B. & Neefjes, J. (2018) Mechanisms of lysosomal positioning and movement, Traffic. 19, 761–769.

16. Pu, J., Schindler, C., Jia, R., Jarnik, M., Backlund, P. & Bonifacino, J. S. (2015) BORC, a multisubunit complex that regulates lysosome positioning, Dev Cell. 33, 176–88.

17. Bagshaw, R. D., Callahan, J. W. & Mahuran, D. J. (2006) The Arf-family protein, Arl8b, is involved in the spatial distribution of lysosomes, Biochemical and Biophysical Research Communications. 344, 1186–1191.

18. Hofmann, I. & Munro, S. (2006) An N-terminally acetylated Arf-like GTPase is localised to lysosomes and affects their motility, J Cell Sci. 119, 1494–503.

19. Qiu, X., Li, Y., Wang, Y., Gong, X., Wang, Y. & Pan, L. (2023) Mechanistic Insights into the Interactions of Arl8b with the RUN Domains of PLEKHM1 and SKIP, J Mol Biol. 435, 168293.

20. Menon, D., Bhapkar, A., Manchandia, B., Charak, G., Rathore, S., Jha, R. M., Nahak, A., Mondal, M., Omrane, M., Bhaskar, A. K., Thukral, L., Thiam, A. R. & Gandotra, S. (2023) ARL8B mediates lipid droplet contact and delivery to lysosomes for lipid remobilization, Cell Rep. 42, 113203.

21. Cabana, V. C. & Lussier, M. P. (2022) From Drosophila to Human: Biological Function of E3 Ligase Godzilla and Its Role in Disease, Cells. 11.

22. Deshar, R., Moon, S., Yoo, W., Cho, E. B., Yoon, S. K. & Yoon, J. B. (2016) RNF167 targets Arl8B for degradation to regulate lysosome positioning and endocytic trafficking, Febs j. 283, 4583–4599.

23. Deshaies, R. J. & Joazeiro, C. A. (2009) RING domain E3 ubiquitin ligases, Annual review of biochemistry. 78, 399–434.

24. Yang, Q., Zhao, J., Chen, D. & Wang, Y. (2021) E3 ubiquitin ligases: styles, structures and functions, Molecular Biomedicine. 2, 23.

25. Buetow, L. & Huang, D. T. (2016) Structural insights into the catalysis and regulation of E3 ubiquitin ligases, Nat Rev Mol Cell Biol. 17, 626–42.

26. Roberts, C. G., Kaur, S., Ogden, A. J., Divine, M. E., Warren, G. D., Kang, D., Kirienko, N. V., Geurink, P. P., Mulder, M. P. C., Nakayasu, E. S., McDermott, J. E., Adkins, J. N., Aballay, A. & Pruneda, J. N. (2024) A functional screen for ubiquitin regulation identifies an E3 ligase secreted by <PSEUDOMONAS aeruginosa>, bioRxiv, 2024.09.18.613774.

27. van Dijk, J. R., Yamazaki, Y. & Palmer, R. H. (2014) Tumour-associated mutations of PA-TM-RING ubiquitin ligases RNF167/RNF13 identify the PA domain as a determinant for endosomal localization, Biochem J. 459, 27–36.

28. Cabana, V. C., Bouchard, A. Y., Sénécal, A. M., Ghilarducci, K., Kourrich, S., Cappadocia, L. & Lussier, M. P. (2021) RNF13 Dileucine Motif Variants L311S and L312P Interfere with Endosomal Localization and AP-3 Complex Association, Cells. 10.

29. Liu, W., Wang, Y., Liu, S., Zhang, X., Cao, X. & Jiang, M. (2024) E3 Ubiquitin Ligase RNF13 Suppresses TLR Lysosomal Degradation by Promoting LAMP-1 Proteasomal Degradation, Advanced Science, 2309560.

30. Zhang, Q., Li, Y., Zhang, L., Yang, N., Meng, J., Zuo, P., Zhang, Y., Chen, J., Wang, L., Gao, X. & Zhu, D. (2013) E3 ubiquitin ligase RNF13 involves spatial learning and assembly of the SNARE complex, Cell Mol Life Sci. 70, 153–65.

31. Trinh, L. T., Van Vu, A., Shin, S., Lee, C., Park, S. H., Cho, E. B., Heo, Y., Kim, H. J., Kim, S. & Yoon, J. B. (2025) RNF13 mediates pH- and Ca(2+)-dependent regulation of lysosomal positioning, Cell Rep. 44, 116053.

32. Yau, R. & Rape, M. (2016) The increasing complexity of the ubiquitin code, Nat Cell Biol. 18, 579–86.

33. Khatter, D., Sindhwani, A. & Sharma, M. (2015) Arf-like GTPase Arl8: Moving from the periphery to the center of lysosomal biology, Cell Logist. 5, e1086501.

34. Jongsma, M. L., Bakker, J., Cabukusta, B., Liv, N., van Elsland, D., Fermie, J., Akkermans, J. L., Kuijl, C., van der Zanden, S. Y., Janssen, L., Hoogzaad, D., van der Kant, R., Wijdeven, R. H., Klumperman, J., Berlin, I. & Neefjes, J. (2020) SKIP-HOPS recruits TBC1D15 for a Rab7-to- Arl8b identity switch to control late endosome transport, Embo j. 39, e102301.

35. Dominska, M. & Dykxhoorn, D. M. (2010) Breaking down the barriers: siRNA delivery and endosome escape, Journal of Cell Science. 123, 1183–1189.

36. Johnson, D. E., Ostrowski, P., Jaumouillé, V. & Grinstein, S. (2016) The position of lysosomes within the cell determines their luminal pH, J Cell Biol. 212, 677–92.

37. Pu, J., Guardia, C. M., Keren-Kaplan, T. & Bonifacino, J. S. (2016) Mechanisms and functions of lysosome positioning, Journal of Cell Science. 129, 4329–4339.

38. Rawat, S., Chatterjee, D., Marwaha, R., Charak, G., Kumar, G., Shaw, S., Khatter, D., Sharma, S., de Heus, C., Liv, N., Klumperman, J., Tuli, A. & Sharma, M. (2023) RUFY1 binds Arl8b and mediates endosome-to-TGN CI-M6PR retrieval for cargo sorting to lysosomes, J Cell Biol. 222.

39. Cantalupo, G., Alifano, P., Roberti, V., Bruni, C. B. & Bucci, C. (2001) Rab-interacting lysosomal protein (RILP): the Rab7 effector required for transport to lysosomes, Embo j. 20, 683–93.

40. Cabana, V. C., Bouchard, A. Y., Sénécal, A. M., Cappadocia, L. & Lussier, M. P. (2025) RNF13 is a novel interactor of iduronate 2-sulfatase that modifies its glycosylation and maturation, bioRxiv, 2025.06.20.660705.

41. Ghilarducci, K., Cabana, V. C., Harake, A., Cappadocia, L. & Lussier, M. P. (2022) Membrane Targeting and GTPase Activity of Rab7 Are Required for Its Ubiquitination by RNF167, Int J Mol Sci. 23.

42. Luzio, J. P., Parkinson, Michael D. J., Gray, Sally R. & Bright, Nicholas A. (2009) The delivery of endocytosed cargo to lysosomes, Biochemical Society Transactions. 37, 1019–1021.

43. Van Wesenbeeck, L., Odgren, P. R., Coxon, F. P., Frattini, A., Moens, P., Perdu, B., MacKay, C. A., Van Hul, E., Timmermans, J. P., Vanhoenacker, F., Jacobs, R., Peruzzi, B., Teti, A., Helfrich, M. H., Rogers, M. J., Villa, A. & Van Hul, W. (2007) Involvement of PLEKHM1 in osteoclastic vesicular transport and osteopetrosis in incisors absent rats and humans, J Clin Invest. 117, 919–30.

44. Lussier, M. P., Cayouette, S., Lepage, P. K., Bernier, C. L., Francoeur, N., St-Hilaire, M., Pinard, M. & Boulay, G. (2005) MxA, a member of the dynamin superfamily, interacts with the ankyrin-like repeat domain of TRPC, J Biol Chem. 280, 19393–400.

45. Mirdita, M., Schütze, K., Moriwaki, Y., Heo, L., Ovchinnikov, S. & Steinegger, M. (2022) ColabFold: making protein folding accessible to all, Nature Methods. 19, 679–682.

46. Abramson, J., Adler, J., Dunger, J., Evans, R., Green, T., Pritzel, A., Ronneberger, O., Willmore, L., Ballard, A. J., Bambrick, J., Bodenstein, S. W., Evans, D. A., Hung, C.-C., O’Neill, M., Reiman, D., Tunyasuvunakool, K., Wu, Z., Žemgulytė, A., Arvaniti, E., Beattie, C., Bertolli, O., Bridgland, A., Cherepanov, A., Congreve, M., Cowen-Rivers, A. I., Cowie, A., Figurnov, M., Fuchs, F. B., Gladman, H., Jain, R., Khan, Y. A., Low, C. M. R., Perlin, K., Potapenko, A., Savy, P., Singh, S., Stecula, A., Thillaisundaram, A., Tong, C., Yakneen, S., Zhong, E. D., Zielinski, M., Žídek, A., Bapst, V., Kohli, P., Jaderberg, M., Hassabis, D. & Jumper, J. M. (2024) Accurate structure prediction of biomolecular interactions with AlphaFold 3, Nature. 630, 493–500.

47. Tauriello, G., Waterhouse, A. M., Haas, J., Behringer, D., Bienert, S., Garello, T. & Schwede, T. (2025) ModelArchive: A Deposition Database for Computational Macromolecular Structural Models, Journal of Molecular Biology. 437, 168996.

48. Wishart, D. S., Feunang, Y. D., Marcu, A., Guo, A. C., Liang, K., Vázquez-Fresno, R., Sajed, T., Johnson, D., Li, C., Karu, N., Sayeeda, Z., Lo, E., Assempour, N., Berjanskii, M., Singhal, S., Arndt, D., Liang, Y., Badran, H., Grant, J., Serra-Cayuela, A., Liu, Y., Mandal, R., Neveu, V., Pon, A., Knox, C., Wilson, M., Manach, C. & Scalbert, A. (2018) HMDB 4.0: the human metabolome database for 2018, Nucleic Acids Res. 46, D608–d617.

49. Jumper, J., Evans, R., Pritzel, A., Green, T., Figurnov, M., Ronneberger, O., Tunyasuvunakool, K., Bates, R., Žídek, A., Potapenko, A., Bridgland, A., Meyer, C., Kohl, S. A. A., Ballard, A. J., Cowie, A., Romera-Paredes, B., Nikolov, S., Jain, R., Adler, J., Back, T., Petersen, S., Reiman, D., Clancy, E., Zielinski, M., Steinegger, M., Pacholska, M., Berghammer, T., Bodenstein, S., Silver, D., Vinyals, O., Senior, A. W., Kavukcuoglu, K., Kohli, P. & Hassabis, D. (2021) Highly accurate protein structure prediction with AlphaFold, Nature. 596, 583–589.

